# CUR(E)ating a New Approach to Study Fungal Effectors and Enhance Undergraduate Education through Authentic Research

**DOI:** 10.1101/2023.02.14.528535

**Authors:** Gengtan Li, Mai McWilliams, Matheus Rodrigues, Benjamin Mearkle, Nader Jaafar, Vivek Golla, Houlin Yu, He Yang, Dilay Hazal Ayhan, Kelly Allen, Domingo Martínez-Soto, Amy Springer, Li-Jun Ma

**Author notes:** Equal contribution (Gengtan Li, Mai McWilliams). Undergraduate students (Mai McWilliams, Matheus Rodrigues, Benjamin Mearkle, Nader Jaafar, Vivek Golla). Peking University Institute of Advanced Agricultural Sciences, Weifang, Shandong 261000, China (Dilay Hazal Ayhan).

## Abstract

Course-based Undergraduate Research Experiences (CUREs) integrate active, discovery-based learning into undergraduate curriculums, adding tremendous value to Biochemistry and Molecular Biology (BMB) education. There are multiple challenges in transforming a research project into a CURE, such as the readiness of students, the time commitment of the instructor, and the productivity of the research. In this article, we report a CURE course developed and offered in the University of Massachusetts Amherst BMB Department since 2018 that addresses these challenges. Our CURE focuses on fungal effectors which are proteins secreted by a destructive pathogenic fungus *Fusarium oxysporum*, one of the top five most devastating plant pathogens. By studying this group of proteins, students are connected to real-world problems and participate in the search for potential solutions. A three-week “standard Bootcamp” is implemented to help students familiarize themselves with all basic techniques and boost their confidence. Next, molecular cloning, a versatile technique with modularity and repeatability, is used as the bedrock of the course. Our past five years of experience have confirmed that we have developed a novel and feasible CURE protocol. Measurable progress documented by students who took this course includes stimulated active learning and increased career trajectory to pursue hypothesis-based research to address societal needs. In addition, data generated through the course advance ongoing lab research. Collectively, we encourage the implementation of CURE among research-intensive faculty to provide a more inclusive research experience to all students, an important element in predicting career success.

## INTRODUCTION

Studying the molecular mechanisms governing all biological systems, the field of biochemistry and molecular biology (BMB) address diverse societal needs, ranging from understanding plant and crop health to diagnosing and treating human diseases. Given this wide range, it is imperative that BMB education keeps pace with ever-increasing scientific innovations, fosters independent and critical thinking, and adequately prepares students to serve the needs of our society. Authentic research experiences add tremendous value to BMB education [1–3] and have a significant impact among women, minorities, and students with disabilities [4]. However, not enough research positions are available to students even at large research-intensive universities. To make authentic research experience available to all, educators are encouraged to replace traditional, cookbook-based laboratory courses with Course-Based Undergraduate Research Experiences (CUREs) by incorporating evidence-based education with enhanced scientific authenticity [5–7]. Enabling a deep connection of material learned in class with the real-world application through a research question, the CURE format offers measurable benefits to students and assists their career development [8]. More importantly, these courses allow undergraduate research experiences to benefit all students and make education more equitable [4,9,10]. In doing so, each lab course is equipped to train students with skills in knowledge discovery, problem- solving, critical thinking, data processing, and collaboration – useful tools for all career paths for all BMB majors and tools that have been proven to better prepare students for life after their undergraduate education [1,11].

While offering a great advantage over traditional laboratory courses, there are multiple challenges at the intersection of a research lab and a teaching lab that manifest when transforming authentic research into a CURE, which explains the fact that the implementation of CURE has been slow, despite realized benefits [12,13]. Due to the challenging pace of research, 13 or 26 weeks can be too short to gather background knowledge, propose hypotheses, acquire research techniques, perform hands-on experiments, troubleshoot, analyze data, and draw conclusions. Faculty may find it challenging to implement a new curriculum by condensing a research project into a one-semester or one-year lab course. This ties into another challenge with the average CURE curriculum, which is the imbalance of scientific research experience among the students. Some students may have prior research experiences and some students may be unconfident to use simple equipment such as pipetting by themselves. Incorporating authentic research requires instructors to meet individual students where they are and make research palatable for students with different levels of background knowledge.

This paper describes a CURE course that serves as an advanced laboratory course for BMB majors, in conjunction with ongoing faculty research. Since 2018, this CURE course has been offered to about 100 undergraduate students in the BMB department at the University of Massachusetts-Amherst. Through this course, students gather data, making contributions to a National Science Foundation-funded research project, “Understanding effector biology in the species complex *Fusarium oxysporum*”. In collaboration with senior scientists in the research lab, students are organized as teams to clone effectors, proteins secreted by the pathogen to manipulate host immunity and attenuate host defense. *F. oxysporum* is a fungal pathogen responsible for destructive and intractable wilt diseases across a diverse spectrum of hosts, including numerous economically important crops, such as banana, cotton, canola, melon, and tomato [14,15]. Research using this model system links the classroom directly with global food security.

In addition to fulfilling a course requirement for our majors, this CURE also helps to boost students’ confidence and provides our majors entering the workforce with critical thinking, troubleshooting, and problem-solving skills. Equally important, students generated valuable materials, including a dozen fungal transformants, each with one effector tagged with a florescent protein, and 100 plasmids that can be expressed either in a host cell or a fungal cell to study effector function and localization. We hope our experience will encourage more scientists to bring their exciting research to the teaching laboratory.

## METHODS

This CURE course (13 weeks, accounting for 4 upper-level credits) was developed in 2017 and has been offered since 2018 as a lab course requirement for BMB Undergraduate Majors. The primary pedagogical goals of this course were to promote student learning in key biochemistry concepts and their application in experimental projects, to work effectively and safely in a biochemistry lab in both individual and collaborative contexts, to establish habits of collaboration and integration with each other, and to improve presentation skills and scientific writing abilities. With these goals in mind, this article highlights how we designed the course to address multiple challenges that faculty encountered when introducing a research topic into a teaching lab. The modularity of the course makes it possible to break the research into three manageable phases.

### Phase I: Prepare Students’ readiness through Bootcamp

A 3-week intensive Bootcamp, developed by the BMB faculty (Supplementary document A), was designed to prepare students both technically and scientifically [16,17]. During the first three weeks of the semester, students were instructed to follow lab safety protocols and conduct fundamental biochemistry and molecular biology experiments, such as measuring the growth curve of the bacteria, bacterial transformation, plasmid extraction (Miniprep), restriction enzyme digestion, PCR amplification, protein induction, purification and quantification, and western blot using standard lab equipment.

As a critical part of the course’ s success, students were introduced to the course’ s research topic through a rigorous research seminar by the course instructor and guest lecturers who are experts in the field. Over the past five years, this CURE module has engaged professors from other institutions, as well as graduate students and postdoctoral scientists from the instructor’ s research team. In addition, students read both review articles and primary publications related to the research topics in a journal club style. By the end of Phase I, each student developed a hypothesis-driven research project by integrating literature knowledge into preliminary data for the list of the proteins provided by the instructor, such as functional prediction, domain similarity, and transcriptional expression.

Throughout the whole semester, students worked in teams formed, with the intent of ensuring diversity among students’ general cultural backgrounds and levels of research experiences. Students within the same team were instructed to rotate roles of leader, recorder, reporter, and skeptic each week. This role rotation ensured students to have equal opportunities, allowed students to respect the effort of the other teammates, and formed healthy and efficient collaboration. A workshop establishing team spirit was implemented in the first week for teammates to get to know each other and establish some ground roles. Each student must form a distinct hypothesis and at the same time, members within each team were encouraged to consult with each other to form a team-based hypothesis.

### Phase II: Perform authentic research using molecular cloning techniques

Students were guided through essential steps involved in cloning a specific effector of interest throughout the course (Figure 1). To develop problem-solving skills and to grow confidence associated with the design, students were given the freedom to repeat experiments and encouraged to schedule meetings with teaching assistants or the instructor to resolve problems derived from each experiment.

**Figure 1.**
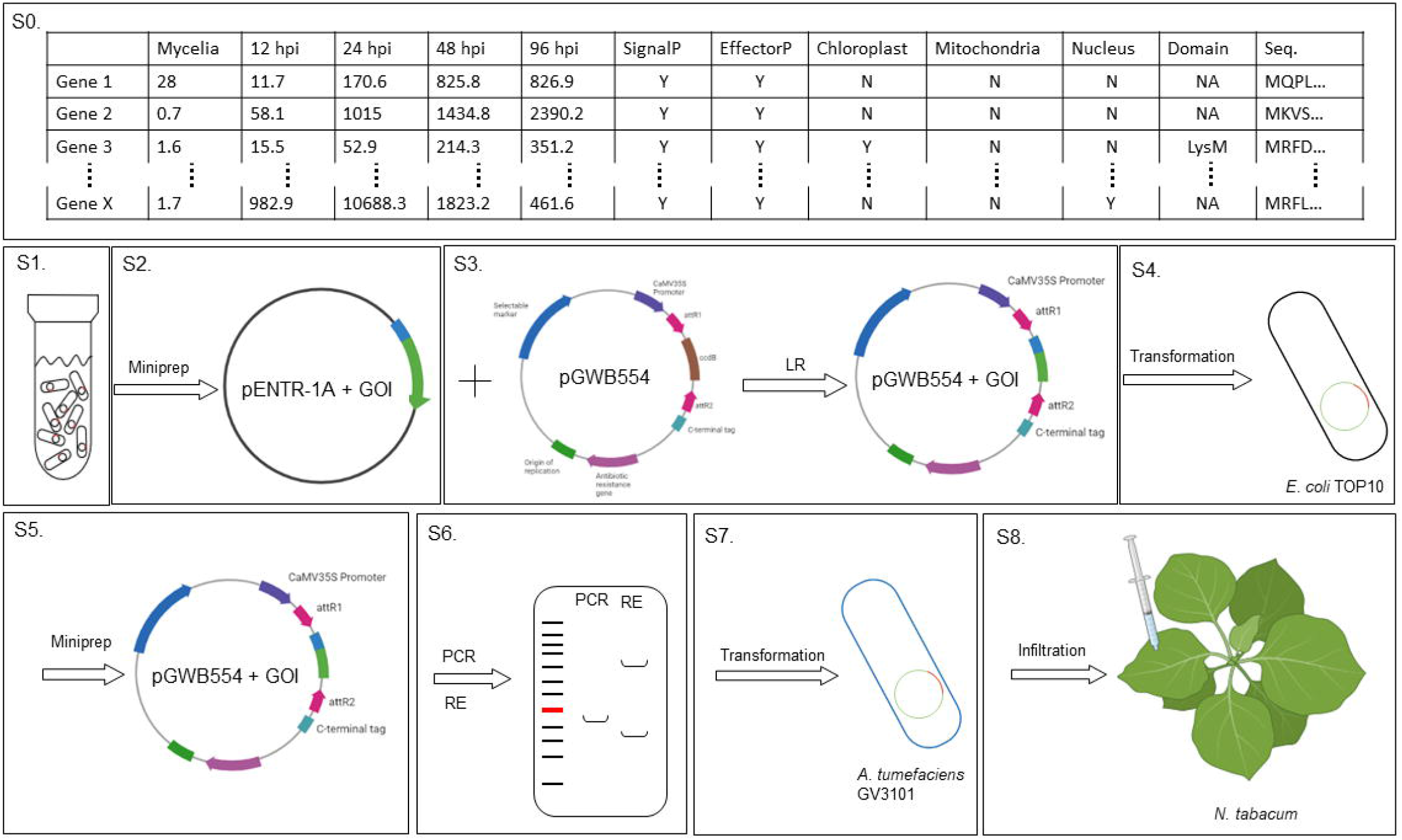
Specific steps involved in generating heterologous expression in *Nicotiana tabacum*. S0. Candidate effector selection. S1. Entry vector preparation. S2. Entry vector extraction. S3. Expression vector generation. S4. Expression vector transformation. S5. Expression vector plasmid amplification and extraction. S6. Expression vector validation. S7. Agrobacterium transformation. S8. Agrobacterium mediated heterologous expression of GOI in *Nicotiana tabacum*.

#### Specific steps

S0. Candidate effector selection: A list of candidate effectors, including the full-length sequence, prediction results, and expression pattern of each effector, was presented to students at the beginning of the semester. Students were guided through protein structure prediction, functional domain recognition, and some relevant references to generate hypotheses.

S1. Entry vector in *E. coli* stock preparation: Entry vectors that contained selected effectors (defined as gene of interests hereafteer, GOIs) were provided to students as *Escherichia coli* stock in week 4. Students reinoculated the bacteria on plates following the Bootcamp bacterial growing protocol. These plates were maintained at 4°C for future use.

S2. Entry vector Extraction: Students extracted plasmid DNA containing the entry vector using the Miniprep protocol introduced in Bootcamp (Supplementary document A).

S3. Expression vector generation: Following the Gateway cloning protocol for LR reaction (Supplementary document B), students made a mixture of the entry vector plasmid with the selected destination vector that contained the Red Fluorescent Protein (RFP) for visualization in a specific ratio. The course TA performed LR reactions for students in the instructor’ s research lab.

S4. Expression vector transformation: The mixture containing the expression vector generated after the Gateway cloning LR reaction was used to transform *E. coli* Top10 cells by following the Bootcamp transformation protocol (Supplementary document A).

S5. Expression vector plasmid amplification and extraction: Students extracted the expression vector from the transformants following the Miniprep protocol.

S6. Expression vector validation: Students performed PCR and Restriction Enzyme digestion following the Bootcamp protocols to check the correct insertion of their GOIs.

S7. Agrobacterium transformation: Once they had confirmed the correct insertion of their effector sequence, students transformed the expression vector into *Agrobacterium tumefaciens* GV3101 following the Agro-transformation protocol (Supplementary document B), and colony PCR was used to confirm the presence of the expression vector in *A. tumefaciens*.

S8. *A. tumefaciens* mediated heterologous expression of GOI in *Nicotiana tabacum*: Students grew the transformants in S7, transferred the *A. tumefaciens* into the infiltration buffer, and infiltrated the 6-week-old *N. tabacum* (Protocol described in Supplementary document B). The expression of GOIs, marked by the RFP signal from the infiltrated plants, were observed using a confocal microscope under the guidance of a graduate teaching assistant.

The cloud-based online notebook Benchling [18] was used to record and organize laboratory observations. Even though research was conducted as a team, we encouraged each student to form the good habit of recording all data individually. The notetaking consisted of three parts: pre-designing the experiment, recording raw data, and completing notes based on data analysis and troubleshooting. Before each lab session, students were asked to write down experimental designs and refer any protocols that was involved in the experiment. During the lab session, raw data was written down informally and after the lab session, students finished their notes by organizing and analyzing raw data, recording modifications of the protocol, formulating conclusions, and documenting errors that occurred during the lab period.

### Phase III: Continue the research through collaboration with the instructor

Once GOIs were cloned, the functional characterization was continued through a collaboration with a senior scientist who was working on the effector in the research lab. One scientist served as a mentor to each team and would continued investigating these GOIs after the completion of the 13-week course in the research laboratory. Results from this CURE course further advanced research prospects in the instructor’ s lab. The results from at least one effector from the previous CURE courses would be used for publication in the coming year and the students who involved in the research would be included as co-authors for the manuscript.

## RESULTS AND DISCUSSION

### The Bootcamp acclimates students before embarking on independent research

Teams were formed with the intent of ensuring diversity among students’ general cultural backgrounds and levels of research experiences. In addition, a workshop implemented in the first week to establish team spirit and foster collaboration among team members had proven to be a crucial component of this preparation phase of the CURE [19]. Team members not only helped each other complete experiments, but also assisted in the growth and development of each other emotionally, scientifically, and technically. As teamwork has become essential for today’ s scientific discoveries, the importance of creating collaborative team spirits cannot be over- emphasized.

Our Bootcamp protocols (Supplementary document A) during Phase I provided step-by-step instructions for all basic analytical and technical skills essential in BMB curriculums, including bacterial transformation, plasmid extraction, polymerase chain reaction, protein induction and purification, and gel electrophoresis. This phase offered students who were not quite comfortable with lab techniques and equipment to grow confidence by learning from their mistakes. This phase also provided students who were experienced in conducting lab research to develop leadership skills by helping others through teamwork and troubleshooting. The Bootcamp experience removed the fear that prevented students from trying. It also enabled instructors to work closely with students to assist them with the obstacles that arise. Finally, the establishment of problem-solving skills increased innovation during the independent research period and set up a strong foundation for the rest of the course.

The other important function of Phase I was to prepare students for an independent investigation. Naturally, students were motivated to do what they decide to do, in contrast to what they were told to do. It is important to introduce essential materials to help students to understand the significance of the research and draw connections to their interests. To introduce students to the research topic, we showed documentaries capturing the devastation of Fusarium wilts and guest speakers working on the fungal effectors to visit the class. Our experiences confirmed that students are very receptive to real-world problems and appreciate scientists who shared authentic research experiences. In addition, to assist in hypothesis generation, students were provided with authentic data and guided through some basic analysis. Datasets include DNA and protein sequences, gene expression patterns, and any additional phenotype the lab had. Basic analyses include BLAST search, protein domain analysis, and structural prediction.

It should be noted that in the past four years, the length of the “Bootcamp” was shortened from 4 weeks to 3 weeks with reduced experiments based on the direct relevance of the CURE research approaches. Thus, the Bootcamp period can be easily implemented and altered depending on the classroom, the topic, and the course length.

### The modularity, flexibility, and multi-channel validation ensure the success of the CURE

During the second phase, students performed authentic research using molecular cloning techniques. We had successfully applied both Gibson Assembly and Gateway cloning techniques, depending on research progress in the instructor’ s lab. Using the Gibson Assembly approach, students designed primers to amplify their GOIs and inserted them into a pre-designed vector prepared by the instructor team. In this case, less preparation was required from the instructor’ s research lab, but the success of this approach required the successful amplification of the GOIs with correct primer design. Using the Gateway cloning approach, each student received an entry vector containing their GOIs. If a library of entry vectors is already prepared in the research lab, this approach offers a higher success rate. Students can select their candidates based on their interests. With multiple destination vectors, students can tag their candidate effectors differently.

For this manuscript, we describe results generated using Gateway cloning (Supplementary document B). We emphasize the possibility of repeating any step, which offers modularity and flexibility to the design. Below are a few critical points for consideration regarding Phase II.

1. The Bootcamp manual offered critical preparation. With experience gained through Bootcamp, the initial steps of conducting original research, from growing *E. coli* stocks with the gene of interest entry vectors to collecting plasmid DNA of the entry vectors and generating expression plasmids through the Gateway cloning protocol, were quite successful.
2. Variability was expected and flexibility was important for all steps after S3. To make the whole class manageable, students were instructed to progress as a team with the freedom to repeat any step through troubleshooting within the team and consultation with the instructor.
3. To build confidence for students to move forward with the procedure, it is important to employ multiple approaches to validate the success of each step. For instance, to confirm the success of transformation, we required two lines of evidence and had implemented 1) plates with a selective medium that confirmed the integration of plasmid with the correct antibiotic resistance gene; 2) colony PCR with specific primers targeting the GOIs; 3) harvest plasmid after the transformation to be subjected for PCR; 4) restriction enzyme digest and/or sequence the harvested plasmid.
4. Observing expressed effectors using successfully transformed expression vectors was the most exciting element at the class-wide level. The *A. tumefaciens* mediated infiltration assay using tobacco plants is a standardized protocol to heterologously express fungal proteins for the study of fungal-plant interactions. Fluorescent protein tags enable the visualization of expressed protein under the microscope (Figure 2C).

**Figure 2.**
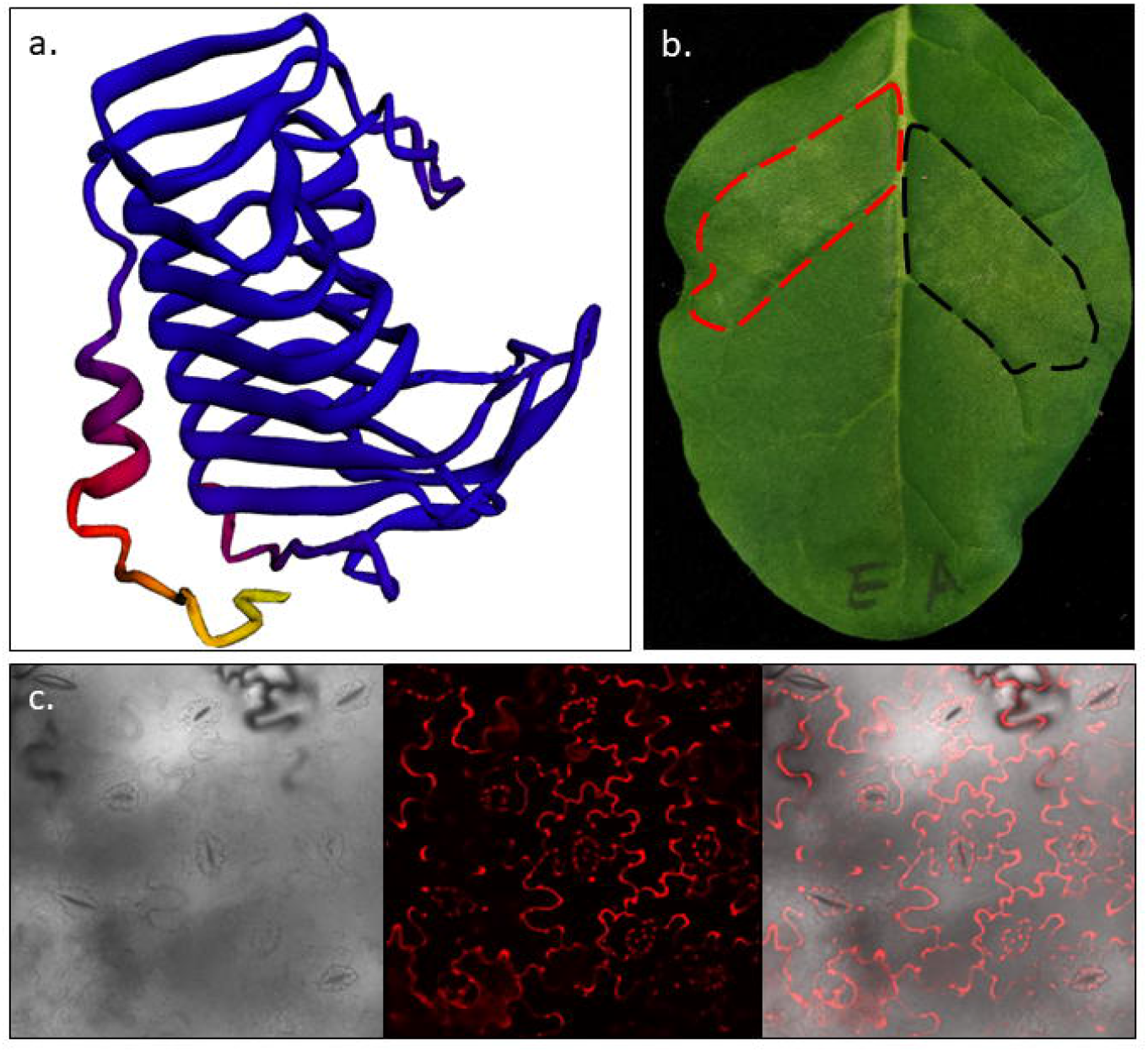
Structure and heterologous expression of effector FOXB_10264 a). Structural prediction of the effector (Gene id: FOXB_10264), signal peptide removed. b). Whole leaf image 3 days after infiltrated with the effector. Infiltration regions are indicated by the dotted lines. Red: Fo5176_411481. Black: Empty vector Agrobacterium. c). Confocal images of 3 days post infiltrated leaf tissue with the effector. Left: Brightfield. Middle: RFP. Right: Merge.

We consider multi-channel validation to be a crucial element of the design. PCR remains a crucial analytical tool for students. However, failure of PCR amplification can be attributed to different technical errors rather than unsuccessful transformation. Repeating PCR amplifications, as well as continuing the validation using alternative approaches after Miniprep, are critical to the speed and success of this second phase of the class.

Individual Benchling notebooks were evaluated every week to recognize successfully performed experiments by each team and to identify common issues. A weekly lab meeting was organized to recognize successful development and dissect challenging issues. Notebook entries were required to include discussion and reflection with the purpose of promoting independence in troubleshooting and problem-solving. Grades were not primarily based on results, but on students’ development and the progression of the project.

### The CURE engages students with excitement and creativity

Most undergraduate students have not experienced any research in fungal effector biology. However, they are smart scientists capable of critical thinking. Once students understand the data they are generating, they can make distinct contributions to the research in their ways. Students with different backgrounds build their hypotheses from a unique point of view. For instance, one team decided to study a group of effectors with plant cell wall degradation activities to test the hypothesis that the secretion of this group of effectors would damage plant cell walls to various degrees. The other team formed a comparative angle to investigate the same group of effectors in and out of the plant cell by including or removing the signal peptide from the candidate effector. Our past experiences had proven that granting students space and freedom encouraged creativity. Over the past 4 years, students had generated valuable materials including a dozen fungal transformants, each with one effector tagged with a florescent protein, and 100 plasmids that can be expressed either in a host cell or a fungal cell to study effector function and localization.

Unlike traditional laboratory courses, the results of the CURE portion were not uniform. To advance the research, each team was assigned to a senior member of the instructor’ s team for further functional analyses. From a research perspective, these mentors shared students’ interests and invested in these effectors as part of their research questions. From a teaching perspective, this arrangement ensured that teams with substantial advanced progress received enough attention for their specific effectors, and at the same time, students who experienced challenges in the early steps would receive sufficient assistance from the instructor and teaching assistants.

Figure 2 illustrates the successful cloning of an apoplastic effector pectin lyase (FOXB_10264) in *F. oxysporum* 5176 [20]. The structural model of the mature protein, generated from RoseTTAFold by removing the signal peptide region [21,22], has a typical pectin lyase fold with a single-stranded, right-handed parallel β-helix topology (Figure 2A), which was highly similar to whole protein structure prediction on AlphaFold [23,24]. Based on the previously reported plant-fungal interaction data, this effector is involved in the course of infection [20] (Figure 2B). The heterologous expression system, where the leaves of *N. tabacum* were infiltrated with *A. tumefaciens* transformed with the RFP-tagged effector, confirms the expression of the effector and illustrates how the effector’ s localization is limited to the apoplast space (Figure 2C). The localization of the effector and its interaction with *N. tabacum*, elucidated with the *A. tumefaciens* mediated infiltration, produced exciting results that could provide insight into the pathogenicity of *F. oxysporum*. Follow-up research on this effector will be included in a manuscript to be submitted within a year, and the student who generated the clone will be included as a co-author of this manuscript.

## CONCLUSION/FUTURE DIRECTIONS

As an inclusive research experience has been accepted as an important element in predicting career success for our undergraduate student [1–3], Course-based Undergraduate Research Experiences (CUREs) add tremendous value to BMB education [3,25]. Here, we describe a CURE module using the molecular cloning technique. This commonly used BMB technique is repeatable and can be standalone or integrated with more functional characterization assays, making the design versatile and adaptable. Equally important, the choice of protein links the bench to the world. Different research teams can invite students to join their research mission by selecting proteins derived from the instructor’ s research. This selection process helps students to establish a connection with real-world topics, creates ownership over the research, and prompts independent thinking. Studying *F. oxysporum* effectors and their roles in plant-fungal interaction brings global food security issues to the classroom. Students are inspired by the possibility of controlling disease through basic research. One of the first authors, Gengtan Li, who took Biochemistry 426 in fall 2019, decided to continue working with the research team and is currently working in the instructor’ s lab as a Ph.D. student.

Students responded positively to this course and reflected increased attraction to science and research, be better prepared for being independent scientists, and increased confidence in conducting research. An essential aspect of a successful scientist is the ability to think critically. Researchers must be capable of overcoming obstacles that bar them from progressing with their work. Our proposed CURE protocol allows undergraduate students to think for themselves rather than following an experimental design with a defined outcome. In the weeks following the Bootcamp, students had to analyze their data and try to understand what they were or were not seeing. Before their effector research, the students were required to read scientific literature on the topic and gain a solid understanding of the research they would pursue for the rest of the semester. Combining both the background knowledge and the result from the research requires attention to detail and extensive analysis to comprehend how the data collected relates to the topic at hand.

Here are a few lessons we learned along the way. First, the success of the design requires preparation before the beginning of the semester, including identifying a list of candidate proteins and testing all protocols. In addition, it is important to emphasize the team-based aspect of the course. Organizing teams with diverse social backgrounds and different research experiences is very important. Intensive preparation at the beginning of the semester is also crucial to help each team to form good work habits and dependable relationships among team members set up a strong foundation for the success of the whole course. Team conferences with the instructor throughout the course also help identify issues and resolve them promptly. The third lesson we learned is the importance of flexibility when implementing the CURE. This is an undergraduate course and mistakes are expected. By granting students the freedom to repeat experiments and identify and correct mistakes, we honor their learning processes. It is also important to implement different means to confirm results.

There is no question that developing and implementing a CURE course is time-consuming, especially during the developmental stages. However, the benefits of doing so are numerous, first and foremost for our students. Traditional lab courses provide students with hands-on experience according to protocols. Taking control of experiments to test hypotheses is difficult in a traditional lab course setting. CURE curricula emphasize independent research, foster scientific inquiry, and assist in the development of future scientists.

Furthermore, we argue that this protocol uniquely prepares students for their future endeavors. Independent research helps students acquire skills in personal-professional communication, organization, collaboration, and independence, all of which are necessary tools in preparing to enter into the workforce [26]. As graduate programs typically prioritize students who have authentic research experience, by implementing such experiences into undergraduate curricula [12], CURE experiences help students continue higher education through graduate school or medical school [27].

A CURE course also offers an equitable approach to bringing research experiences to all BMB majors, rather than solely to the handful who are comfortable reaching out to professors to secure a position in a research lab [9,10]. Research experiences uniquely benefit students who would traditionally perform poorly in lecture-style courses, more so than better-performing students. Similarly, a more uniform research approach that is embedded into the curriculum may help disadvantaged groups by opening access to research opportunities. As both Ph.D. programs and medical schools emphasize the value of research when selecting their top applicants [12], research experience offered through CURE makes education more equitable for all students and better prepares our students for their future professional careers.

Future work should focus on adapting this protocol to different areas of research. The Bootcamp manual serves as an educational tool, incorporating a standard aspect of this curriculum. Faculty interested in implementing this type of curriculum could focus on their expertise and research when students are writing their independent research proposals. Not only can this class help students gain experience conducting research, but it may also introduce students to new research interests and fields of science not taught in the classroom. Additionally, this CURE protocol aids faculty who participate in original research with the unique means of gathering large amounts of data, which benefits faculty research.

## ACKNOWLEDGMENTS

We appreciate UMass Amherst BMB teaching staff, Pa Tamba Ngom, Erica Light, Rachel Cole, Tien Bui, and Nancy Robbins Thorne-Thomsen for their excellent support, for every BMB undergraduate student for supporting the progress of this course, and for the BMB writing fellow Dr. Robin Garabedian for assisting the development of this manuscript. This project is supported by the Natural Science Foundation CAREER award IOS-165241 to L.J.M and NSF-RCN-UBE 2119918 to AS. L.-J.M. is also supported by the National Eye Institute of the National Institutes of Health under award number: R01EY030150, USDA/SCRI/NIFA award (2016-68004-24931), and the USDA National Institute of Food and Agriculture (MAS00496). The funding bodies played no role in the design of the study and collection, analysis, and interpretation of data and the writing of the manuscript.

## AUTHOR CONTRIBUTIONS

This paper was developed in a writing course, Biochem430, through a collaboration among all authors, including students of the writing course (MM, MR, BM, NJ), students of the CURE course Biochem426 (DL, MM, VG) and previous graduate TAs of the CURE course (GL, KA, DHA, HY). L.-J.M. was the instructor for both courses. The co-first authors, GL and MM, lead the writing efforts and compiled all figures. VG cloned Pectin Lyase, used as an example in Figure 2. KA, HY and DMS, senior scientists from the Ma lab, assisted advanced research components. AS lead the development of the bootcamp manual. All authors contributed to the writing.

## SUPPLEMENTARY INFORMATION

Supplementary document A: Bootcamp protocol. Supplementary document B: Cure Methods.

