## Supplementary document 1 for "CUR(E)ating a New Approach to Study Fungal Effectors and Enhance Undergraduate Education through Authentic Research"

### BIOCHEM 426 BOOT CAMP MANUAL, FALL 2020

BMB Teaching Labs

JULY 30, 2020

UNIVERSITY OF MASSACHUSETTS, AMHERST (UMASS AMHERST)
661 N Pleasant Street, Amherst MA 01003 (Integrated Sciences Building (ISB))

### Table of Contents

Schedule ... 6
Welcome to Boot Camp ... 6
Safety in Life Science Laboratories ... 7
Lab Safety Summary ... 9
Review of Micro pipetting ... 9
Micropipettors ... 9
When taking up solution into the pipette tip: ... 9
When expelling liquid: ... 9
Multichannel Pipettor ... 10
Background on Malate Dehydrogenase and Expression System ... 10
Why MDH? ... 10
Watermelon gMDH ... 10
Table 1: Watermelon gMDH properties ... 11
Selecting an expression system ... 11
Host cells ... 11
Expression plasmids ... 11
pQE-WgMDH uses the pQE60 expression plasmid ... 12
Figure 1: pQE-60 Vector map (Source: Qiagen.com) ... 13
Strategies in directional cloning of Watermelon gMDH into pQE60 ... 14
Bacterial transformation, selection, and screening ... 14
Host E. coli transformation strain ... 15
Host *E. coli* expression strain ... 15
Web Resources for Biochem 426 Boot Camp ... 15
Day 1- MB: Computer Cloning Exercise ... 17
Objectives ... 17
Procedure ... 17
Using Serial Cloner to make a plasmid ... 17
Build a file of your vector ... 17
How to Clone: get WgMDH into the pQE60 MCS ... 17
Primers available: (corrected) ... 18
Restriction sites ... 18
Pre-Boot Camp Prep (performed by teaching staff) ... 19

Setting up dilutions of OAA: 65
Plan your reactions: 65
Table 7: Worksheet: 66
Turnover number 67
Procedure 68
Running the Reactions 68
**Table 8: Results:** table for calculation of reaction velocities. 69
Construct Michaelis-Menten (M-M) and Lineweaver-Burk (L-B) plots: 69
Using Excel to Plot Enzyme Kinetics and Inhibition Data: 70
The Michaelis-Menten (M-M) Plot 70
The Lineweaver-Burk (L-B) Plot: 70
Experiment II: Vary conditions that affect the Michaelis-Menten parameters of MDH 70
Table 9: Worksheet for Experiment II 71
Discussion Questions: 71

#### Schedule

| Day & Date | | Techniques | |
|---|---|---|---|
| | | **Protein Purification stream** | **Molecular Biology stream** |
| **1** | | | Computer lab, Start overnight cultures for minipreps |
| **2** | | Plot growth curve of *E. coli* | Plasmid prep and digest |
| **3** | | Induction | DNA gel for digests, Transformation |
| **4** | | Lysis, and Purification | Count colonies and streak |
| **5** | | MDH Assay, and Bradford Assay | Start overnight cultures for culture PCR, restreak if necessary |
| **6** | | Calculate specific activity | Setup PCR, check streaked plates |
| **7** | | SDS-PAGE, Transfer, review specific activity calculations | DNA gel for PCR |
| **8** | | Complete Western Blot: Immunodetection, Enzyme kinetics assay, calculate Vmax and Km | |

#### Welcome to Boot Camp

Boot Camp is a 4-week long skill-intensive workshop designed to prepare you for a Classroom Undergraduate Research Experience (CURE). Boot Camp labs are adapted from many previous years of running this course. They are designed to reliably produce the same results each time. The CURE will take place after Boot Camp for the remainder of the semester and will provide you the opportunity to contribute to the body of scientific knowledge. CURE experiments are original and so have not been performed before. Success during the CURE will require critical thinking, creative problem solving, teamwork, and a robust understanding of the biochemistry and molecular biology lab techniques emphasized in the first month of the semester.

Boot Camp has two "streams" or projects that you will perform simultaneously. One involves Molecular Biology (the "MB" stream), working with plasmids and PCR, and one which involves
Protein Purification (the "PP" stream), in which you will express the gene encoding the protein MDH, purify this protein in large amounts, and characterize its enzymatic function. The schedule for Boot Camp is outlined below.

#### Safety in Life Science Laboratories

BMB laboratory courses are designed with safety in mind. All instructors, support staff, and students must take an active role to promote lab safety; this includes wearing appropriate personal protective equipment (PPE) and proper disposal of hazardous material. See below for further instruction. Any student who has safety concerns, a medical condition, or other circumstances which might impact his/her participation in the labs is strongly encouraged to discuss these circumstances with the instructor as soon as possible.

1. Clothing choice is critical for maintaining personal protection. Shoes must be close-toed (no sandals, flip-flops, etc.), and long pants must be worn in the laboratory. Shirts should have sleeves, and no dangling hair or jewelry is allowed.
2. Lab coats must be worn while in the laboratory
3. Eye protection is required in the laboratory at all times. It is especially critical to wear proper eye protection when there is potential for exposure to ultraviolet (UV) light, caustic chemicals, items under pressure, aerosol and where a broken/flying glass hazard exists.
4. Protective gloves of the proper type should be used when necessary.
5. COVID19 safety requirements in the BMB Teaching labs also include wearing a facemask, face shield and to practice social distancing of at least 6 feet apart. All other COVID19 specific protocols provided, including cleaning and disinfection, must be strictly followed.

6. To minimize cross-contamination and distraction, students should not use cell phones or iPods while in the laboratory room.

7. Maintaining a clean, clutter-free working environment promotes safety. All jackets, sweaters, backpacks, etc., as well as any materials not needed in the lab, should be placed in the cubbies at the entryway door. Lab benches should be wiped with 70% ethanol prior to and upon completing experiments.

8. All students should wash their hands with antibacterial soap and warm water upon entering and leaving the laboratory room.

9. The laboratory contains several safety features that each student should note upon first entering the room. Each lab room has a drenching shower, eyewash station, first aid kit, and fire extinguisher.

10. All students should be familiar with the emergency exits (each lab room has two exits) and escape plan from their lab room and all common areas.

11. Never bring food or beverage into the laboratory or consume food or beverage in the laboratory. Additionally, do not dispose of food wrappers or beverage containers in the lab wastebaskets, even if the contents were not consumed in the lab. Inspectors conclude from such evidence that food or drink has been consumed in the lab room. Note that chewing gum counts as food under this rule.

12. All students should be familiar and comfortable with the proper operation of all instrumentation as well as the procedures to be performed in the lab. Students should be aware of the hazards that exist in the laboratory. Do not hesitate to ask the instructor or TAs for help. Report any instrument malfunction immediately to the teaching staff.

13. Label all bottles, flasks, test tubes, etc., that you use. A proper label should contain your name or initials, contents, concentration, course number, and date.

14. Never pipette by mouth. Remove only as much as you need for the experiment using the proper delivery method. Never return any excess to the reagent bottles, as it may cause contamination of a common supply.

15. You are responsible for safe handling and disposal of all chemicals, solutions, and equipment you use. Do not leave chemicals around common areas or your lab bench for labmates or the teaching staff to clean up. Proper disposal of laboratory materials is critical.

a. The large gray trash bins are for paper towels, kimwipes, gloves, tape, and other general Lab trash that is not sharp or contaminated. Do not put liquids, sharps, biohazards, broken glass, or pipets/tips in these trash barrels!

b. Pipet tips and plastic pipets (along with microfuge tubes and empty conical tubes) should be deposited in the plastic beakers located on the benchtop. TAs will properly dispose of these items at the conclusion of each laboratory session.

c. Broken glass must safely be deposited in the "glass waste disposal" cardboard boxes. Do not overfill these containers, and do not put items that do not belong into these containers (including gloves, plastic pipets, general lab trash, and liquids).

d. Sharps such as razor blades and needles must be placed into the red "sharps" container.

e. Each liquid hazardous waste will have a dedicated container in the fume hood for proper disposal. Consult with the teaching staff on the location of these containers.

f. Biohazardous materials or items that require decontamination (i.e., bacterial growth plates/cultures, transgenic plants, etc.) will be collected by the teaching staff.

15. Our laboratory courses are designed with safety in mind. Any student who has safety concerns, a medical condition, or other circumstances which might impact his/her participation in the labs is strongly encouraged to discuss these circumstances with the instructor as soon as possible.

#### Lab Safety Summary

All instructors, support staff, and students must take an active role to promote and maintain a safe laboratory environment. Following the requirements and guidelines provided by the Environmental Health and Safety office (EH&S) and adhering to the above guidelines will maximize safety and enjoyment for everyone!

#### Review of Micro pipetting

##### Micropipettors

Micropipettors are an essential piece of lab equipment. The accuracy of pipetting always determines the success of an experiment. Dropping a micropipettor will bend the piston assembly and ruin its accuracy. Take care of your micropipettors because you will be using the same ones throughout the semester.

Each micropipettor has a range of accuracy (see below). Even though the dial will theoretically allow you to pipette outside of that range, do not do so.

P20: Use for 1 µL to 20 µL
P200: Use for 20 µL to 200 µL
P1000: Use for 200 µL to 1000 µL
Make sure you dial the correct volume!

Never attempt to draw up liquid without a pipette tip firmly attached to the micropipettor. Note that when pressing down the plunger, there are two stops.

##### When taking up solution into the pipette tip:

With the pipette tip outside of the solution, depress to the first stop only, place the tip into the solution then slowly release to draw up liquid (rapid release results in aspiration of liquid into the barrel of the micropipettor, which compromises accuracy). Always check the pipette tip to verify that approximately the correct amount of solution was taken up—you need to develop a sense of what 1 µL or 10 µL looks like in the pipette tip. Check with an instructor for help.

##### When expelling liquid:

Place the liquid-filled tip into the tube, and while touching the inside of tube, depress to the second stop. Surface tension will prevent release of all of the liquid unless the tip is touching the tube. Before releasing, ensure the pipette tip is fully removed from the liquid to prevent reuptake.

When adding solutions to microfuge tubes: always hold the tube tilted at an angle and add solution to the side of the tube near the bottom. Always hold the tube in your other hand. Touch the pipette to the wall of the tube and watch to see that the liquid comes out of the tip.

##### Multichannel Pipettor

The multichannel pipette is a way to transfer multiple small volumes of liquid at once, such as to a 96 well plate. These may be used during some experiments throughout the lab course when necessary. The volume on a multichannel pipette can be adjusted using the arrow keys and confirmed with the enter key. Taking up and expelling liquid are both done by pressing the "pipette key" on the manifold. NOTE: when taking up liquid, keep the pipette tips vertical; when discharging liquid, remove tips from solution immediately after discharge to prevent reuptake of liquid.

#### Background on Malate Dehydrogenase and Expression System

##### Why MDH?

Malate Dehydrogenases catalyze the reaction:  Malate + NAD+ into Oxaloacetate + NADH, involving a simple hydride transfer from the 2 position of Malate to the nicotinamide ring of NAD+ to give NADH. During the process, a proton is also released (Goward and Nichols. Protein Science 1994; 3:1883-1888). This reaction plays a number of roles important to metabolism, the most familiar is the step in the citric acid cycle, but is also critical to the Urea cycle and Gluconeogenesis, and is one of two key enzymes in the Malate-aspartate shuttle that transports reducing equivalents between cytoplasmic and organelle compartments in cells. Modulations of these pathways are critical in the progression of many diseases, including cancer, as well as in the adaptations of pathogens to their human hosts.

There are many malate dehydrogenases- it is a protein family, and members differ slightly in amino acid sequence from one another.  They are all related by evolutionary ancestor, have the same critical residues in their active sites, and carry out the same reaction.  However, one malate dehydrogenase protein might be found only in the mitochondria of one organism and another is found only in the cytoplasm.

##### Watermelon gMDH

Our protein was originally isolated from the glyoxysomes of germinating watermelon seedlings. Glyoxysomes are organelles of plants, involved in various biosynthetic pathways as well as fatty acid oxidation.   We refer to this protein as Watermelon ( *Citrullus lanatis* ) glyoxysomal malate dehydrogenase, or WgMDH.   (For reference, see: Cox et al. FEBS J. 2005; 272:643-654).

#### Table 1: Watermelon gMDH properties

| | |
|---|---|
| Size | 326 residues (978 nucleotide bases) |
| Molecular Weight | ~34.3 kiloDaltons (kD) |
| Isoelectric point (pI) | ~8.3 |

#### Selecting an expression system

Selection of the appropriate host cell system and expression vector is necessary to optimally express the target protein. Cell systems for expressing protein include bacteria, yeast, animal cells, and insect cells. Selection of an expression system is often dependent upon the chemistry and characteristics of the protein you wish to express. Each cell system utilizes unique expression vectors (typically plasmids) to appropriately drive protein synthesis; the expression vector must be compatible with the host cell, and typically possesses desirable features to optimize protein expression and enhance product recovery and purification. Knowing the specific characteristics of the host expression system and the target protein to be expressed is critical in product recovery and purification.

##### Host cells

Bacteria (typically *E. coli)* are nearly always capable of expressing a protein from a sub-cloned gene, and many plasmids are readily available to express the target protein. *E. coli* are unicellular and prokaryotic (lack a membrane-enclosed nucleus), and thus transcription of the gene into RNA and subsequent translation into protein are coupled processes. Proteins expressed in E. *coli can* reside in many different fractions, both intracellular (cytoplasmic) and extracellular (if secreted into the culture medium), and may be present in a soluble or insoluble form.

Many researchers have previously validated that watermelon gMDH can be expressed and recovered in active form from bacteria. This gMDH is expressed from cDNA, generated from mRNA lacking introns (If introns were present in the clone, they could not be removed by E. *coli* and expression would not produce MDH protein).

##### Expression plasmids

In addition to simply being compatible with the chosen host cell, expression plasmids are optimized to (1) synthesize high levels of protein, and (2) aid in recovery and purification of the expressed protein in downstream steps. Expression plasmids contain the following features to ac accomplish the above two goals:

A. An origin of replication (or genome insertion mechanism) to ensure the plasmid is replicated in the host cell.

B. A selectable marker to make sure the cells containing the plasmid are isolated and the plasmid is maintained during host cell culture.

C. A multiple cloning site (MCS) containing multiple unique restriction enzyme sites to allow for insertion of the target gene at a specific location.

D. A host cell-compatible promoter sequence upstream from the target gene to drive protein expression. Transcriptional terminator sequences are sometimes employed to optimally express the proper transcript.

E. An affinity tag, a specific sequence of DNA that encodes an amino acid sequence recognized in a highly specific fashion by a target molecule (i.e., peptides, antibodies, metals, substrates, etc.). The affinity tag is covalently fused to the target protein upon expression. If the target molecule is immobilized on an inert bead, resin, or column, the protein covalently coupled to the tag can be specifically and easily affinity purified. The affinity tag can be placed on either the N- or C-terminus of the target protein, or less commonly within the target protein.

#### pQE-WgMDH uses the pQE60 expression plasmid

We have used an expression plasmid from Qiagen Inc. to express gMDH in E. *coli*. pQE60 uses a promoter sequence derived from the T5 bacteriophage (pT5), followed closely in the sequence by two copies of the lac operator, then a translational start signal (RBS), and ATG start codon. The MCS is followed by sequence encoding a series of 6 histidines (6xHis), to be fused to the expressed protein of interest to form a “His tag.” 6xHis binds with high affinity to nickel ions, a feature that is exploited for purification. pQE60 also expresses the ampicillin resistance ($Ap^r$) selectable marker, relies on the host cell expression of the lac repressor (lacI gene), and uses the Col EI origin of replication.

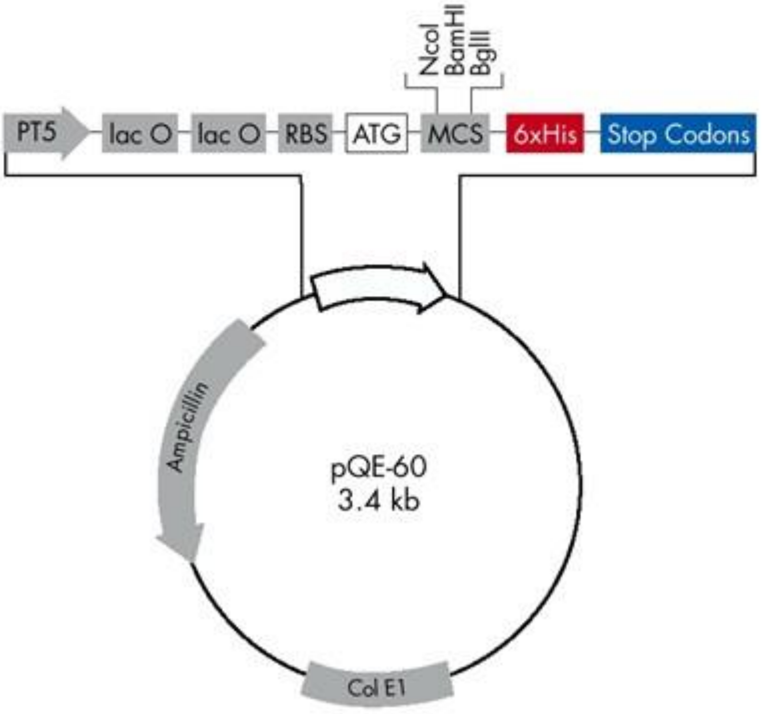

**pQE-60 Vector**

| Positions of elements in bases | |
|---|---|
| Vector size (bp) | 3431 |
| Start of numbering at *XhoI* (CTCGAG) | 1–6 |
| T5 promoter/lac operator element | 7–87 |
| T5 transcription start | 61 |
| 6xHis-tag coding sequence | 133–150 |
| Multiple cloning site | 113–132 |
| Lambda $t_0$ transcriptional termination region | 173–267 |
| *rrnB* T1 transcriptional termination region | 1033–1131 |
| ColE1 origin of replication | 1608 |
| β-lactamase coding sequence | 3226–2366 |

Figure 1: pQE-60 Vector map (Source: Qiagen.com)

The pQE system utilizes elements of the lactose operon to regulate protein expression. Specifically, the engineered promoter contains two copies of the lac operator that is recognized by the lac repressor (lacI) in the absence of lactose. Presence of the non-metabolizable lactose analog IPTG relieves repression at the promoter (LacI protein cannot bind the operators under these conditions), and transcription of the gene(s) under its control is accomplished using the bacterial RNA polymerase. Under ideal conditions, the protein of interest represents as much as 20-40% of total cellular protein, with typical yields in the 10-40 mg per liter of bacterial culture.

Our MDH protein has a tag of 6 histidine residues (6xHis) on its C-terminus. Purification of MDH will utilize the specificity of binding of the His tag to nickel in affinity chromatography.

#### Strategies in directional cloning of Watermelon gMDH into pQE60

In order to assure that the MDH protein would be expressed, the sequence needs to follow in the same direction as the T5 promoter. The restriction sites in the MCS can be used to assure the gene is inserted in the correct direction, by adding an NcoI site at the 5 end of the gMDH gene, and a BglII site at the 3' end, and then opening the pQE60 with these two restriction enzymes, so that the gene can only be inserted in one direction; the Ncol-site also provided the start codon. These restriction sites were added to the MDH sequence by incorporating them into primers used to amplify the gMDH gene using PCR. If the same restriction site had been put on each end of the MDH sequence the gene would have been able to ligate into the plasmid in two equally probable orientations, one of which would be backwards and could not be expressed in this system. Using two different sites confers a directionality to how the target genes are ligated into the plasmid. In addition, careful attention must be paid when designing primer sequences for cloning into affinity tag-containing plasmids, as the target DNA must be inserted in the same reading frame as the tag.

The mature gene sequence without signal sequence was used for this clone (Biochemica et Biophysica Acta 1274 (1996) 48-58; and FEBS Journal 272 (2005) 643-654). Sequences of full-length protein can be found in: UniProt: P19446, Gene: MDHG_CITLA. PDB 1smk

Forward (5') Primer:

MDH NcoI f

GAGTCCATGGCTAAAGGCGG ~Tm= 62ºC

Reverse (3') Primer: MDH Bgl II r

GAGTAGATCTGCTTCTGATG ~Tm= 62ºC

Compare the sequences above with the gMDH gene sequence, and see if you can find the start of the gene, and the end of the gene (is the stop codon included in this primer or not?). Where are the restriction sites in these primers? (start, end or middle of the sequences? Why are they placed thus? How can you tell if the gene will end up in the correct reading frame to incorporate the His tag? These are considerations taken into account when designing primers.

#### Bacterial transformation, selection, and screening

In order to generate large and renewable quantities of our plasmid with inserted target gene, the plasmid is transformed into E. *coli*. To do this, *E. coli* must be made competent to uptake DNA (DNA is a large, negatively charged molecule that must cross the cell membrane); this is accomplished for routine sub cloning by treating with $Ca^{2+}$ ions on ice (high efficiency cloning of libraries may use electroporation to transform DNA). $Ca^{2+}$ treated cells are mixed with DNA to be transformed on ice, then a heat shock step (42ºC) is administered to permit the DNA to cross the cell membrane. Cells are allowed to recover cell wall integrity and express the selectable marker (usually encoding antibiotic resistance) by incubation at 37ºC in medium.

Cells are then plated on the selective medium (antibiotic), upon which only the very few cells that took up the plasmid can grow. This process of isolating only the cells transformed with plasmid is called selection. Screening is the process of identifying those selected cells that contain not just a plasmid, but likely the plasmid of interest. We will screen by using PCR to amplify the MCS of the plasmid. Insertion will generate a PCR fragment much greater than the length of the empty MCS. The putative clones identified in the PCR screen would then be further validated by techniques such as restriction enzyme mapping and DNA sequencing.

##### Host E. coli transformation strain

We perform initial transformations with the E. coli strain DH5α. There are many, many strains of *E. coli* used in molecular biology laboratories, so it is important to always make note of which one you are using. DH5α has been engineered to maximize transformation efficiency. This strain is characterized by three mutations: recA1 inactivates recombination, end A1 prevents degradation of the inserted plasmid, and lacZM15 allows for blue-white screening. Derivatives of DH5α are common and are optimized with additional mutations. A new construct must be transformed into a high efficiency transformation strain before it can be replicated and transformed into an expression strain.

##### Host *E. coli* expression strain

We perform expression in the *E. coli strain* BL21. BL21 has been engineered to express high levels of the target protein in an inducible fashion, and lacks two major bacterial protease genes, lon and ompT, thus reducing the potential for degradation of the highly expressed protein. For our expression and purification experiments, cultures will be induced with IPTG, which will lead to a rapid expression of the MDH-6xHIS protein. The growth medium will be LB supplemented with ampicillin to continually select for plasmid. The pQE-based plasmid uses the host cell's lacI gene, which expresses the lac repressor, essential for minimizing "leaky" expression of MDH in the uninduced state.

#### Web Resources for Biochem 426 Boot Camp

**Watermelon gMDH sequence (protein or DNA):**
http://www.rcsb.org/pdb/explore/remediatedSequence.do?structureId=1SEV

**More information about (and sequence of) the pQE60 plasmid**
https://www.qiagen.com/us/shop/sample-technologies/protein/expression-purification-detection/c-terminus-pqe-vector-set/#productdetails

https://www.addgene.org/vector-database/3881/

**Handbook on protein expression:**

https://www.qiagen.com/us/resources/resourcedetail?id=79ca2f7d-42fe-4d62-8676-4cfa948c9435&lang=en

**Primer3Plus (primer design):**
http://www.bioinformatics.nl/cgi-bin/primer3plus/primer3plus.cgi

**New England Biolabs tools (restriction enzyme and digest information, restriction maps, Tm calculator, plasmid analysis):**
https://www.neb.com/tools-and-resources/interactive-tools

**Dolan DNA Learning Center (animations of molecular biology techniques)**
http://www.dnalc.org/resources/animations/

**Free Plasmid Drawing Software**
Serial cloner: http://serialbasics.free.fr/Serial_Cloner.html
APE http://biologylabs.utah.edu/jorgensen/wayned/ape/
Scalable Vector Graphics
http://www.bioinformatics.org/savvy
Snapgene http://www.snapgene.com
Benchling https://www.benchling.com/molecular-biology/

**National Center for Biotechnology Information (NCBI)**
http://www.ncbi.nlm.nih.gov/.

**Primer Blast**
http://www.ncbi.nlm.nih.gov/tools/primer-blast/

### Day 1- MB: Computer Cloning Exercise

#### Objectives

Today you will use a program called Serial Cloner to step through the process of cloning a gene using the Polymerase Chain Reaction (PCR). You will use the Watermelon glyoxysomal MDH (WgMDH), cloned into the commercial expression plasmid pQE60 as described in the introductory section “Selecting an expression system.”

#### Procedure

You will need to have a computer onto which you can download Serial Cloner (Serialbasics.free.fr). If you do not have a computer to use, you can borrow one in the lab.

##### Using Serial Cloner to make a plasmid

1. Start Serial Cloner (see link above for download)
2. File>New a window will open, paste the sequence of the Watermelon glyoxysomal MDH (WgMDH) gene in, **SAVE THIS.**
3. Repeat with the pQE60 sequence, **SAVE THIS**.

##### Build a file of your vector

4. Continue working with the pQE60 sequence
   Circularize the sequence (in the “sequence” menu, be sure the sequence is unlocked)
   Open the “features” tab. There are no features, but you can add them manually from the image of pQE60
5. Add features to your pQE60
   -In Features window, click on scan to find many standard plasmid features (AmpR, origin, etc.)
   **SAVE** this file
   Now re-open the Graphic map: you should see all the features indicated.
   Restriction sites: Can you find the mcs?
      You can show: unique sites only, sites that occur twice (double) or pick specific sites

#### How to Clone: get WgMDH into the pQE60 MCS

→ Use PCR with primers that create restriction sites (List of primers available is on reverse side)

Note that each primer has (a) homology to part of WgMDH, (b) a restriction site, and (c) a “spacer”, usually about 6 bases, at its 5’ end.

6. Pick 2 primers to use for PCR (from the list below, you can copy and paste these sequences from the electronic version of this file). Hint- make sure the restriction sites will give you a good clone.
   Create a file for each primer. **SAVE** each file.
   For each one, confirm that it matches one end of the WgMDH

How can you do this? Use Find function
Make sure you can recognize: a) which end of WgMDH each primer recognizes, b) what restriction site each has, c) where the "spacer" is

#### Primers available: (corrected)

1. WgMDH BamHI  actagt*ggatcc*atggctaaaggcgg ~ Tm = 59ºC
2. WgMDH EcoRI  ggaggt*gaattc*atggctaaaggcgg ~ Tm = 60ºC
3. WgMDH HindIII  actagt*aagctt*cgaaggaaactcccttctc ~ Tm = 60ºC
4. WgMDH Bgl II  actagt*agatct*gaaggaaactcccttctc ~ Tm = 57ºC
5. WgMDH NcoI  aaaaaa*ccatgg*ctaaaggcgg ~ Tm = 57ºC
6. WgMDH NotI  aaaaaa*gcggccgc*gaaggaaactcccttc ~ Tm = 65ºC

#### Restriction sites

| | | | |
|---|---|---|---|
| BamHI | GGATCC | Bgl II | AGATCT |
| EcoRI | GAATTC | NcoI | CCATGG |
| HindIII | AAGCTT | NotI | GCGGCCGC |

7. Now, create a Virtual PCR product – use the "PCR" toolbar icon
   The part of the primer that matches one end of your gene primer is "segment identical to the target sequence"
   You can use this to add the restriction sites (non-identical sequence) to your WgMDH – as "oligo tail not identical to the target sequence"
   Evaluate and run the PCR. **SAVE** your PCR product as a separate file
8. Use "Construct" to put together your insert (PCR product) and pQE60 vector
   Be careful- you need to have the right insert (with restriction sites as ends, use sticky ends (what are these?) be sure to put the sites into the program in the right order (Might have to try a few times to make it work).

   **To select a site,** click on the restriction enzyme name in the graphic (one click makes it **green** ("start"), two clicks makes is **orange** ("stop"). Note that the "start" in vector is the "stop" in insert. Also, check that you have selected the site in the correct order by noticing how long the pQE60 fragment is after you have done the selection. It should be about 3.4 Kb, if it is only a few bases, you went the wrong way (Might have to try a few times to make it work)

9. Check your ligation result-
   - Are insert and vector portions the right size?
   - Does the gene go in the correct (reading frame) direction?
   - If not- try again
   - If right, SAVE this file.

10. Use “Virtual cut” to verify your plasmid:
   - Find some enzymes that will give you a distinct pattern between your correct ligation, any incorrect ligation, and the pQE60 vector alone.
     **Show your plasmid, and virtual digest, to the instructor**

11. **Check that your fusion protein was made correctly**
Open Graphic Map of your pQEWgMDH construct, so you can see if the construct correctly made a WgMDH- his tag fusion (does the reading frame end in a string of H’s?).
    Use “find”… “ORFs”
    Show the fusion ORF to the instructor
12. Rename the new “feature” that is your MDH-6xHis fusion protein:
    -Open “find” (bottom right of sequence window)
    -select the “ORFs” tab
    - find MDH (what base does it start at?)
    - click on that ORF, it will highlight the DNA sequence in blue, right-click on the highlighted sequence and select “create New Feature from selection”
    - go back to the Features window and rename “new feature” to “MDH”
13. Open graphic map again- note that you have to reopen it every time you change or add a feature. **Show your graphic map to the instructor**

### Pre-Boot Camp Prep (performed by teaching staff)

#### Streaking plasmid pQE_WgMDH for miniprep

On Thursday of the week before Boot Camp starts (Tuesday of following week), DH5α E. coli colonies transformed with the plasmid pQE_WgMDH will be streaked (as per pattern on figure 1 below) onto plates containing LB agar and 100 µg/mL of ampicillin), for the start of over-night cultures on Day 1-PP. These will be used for miniprep of pQE_WgMDH plasmid DNA

Figure 2: Streaking on LB Agar Plate

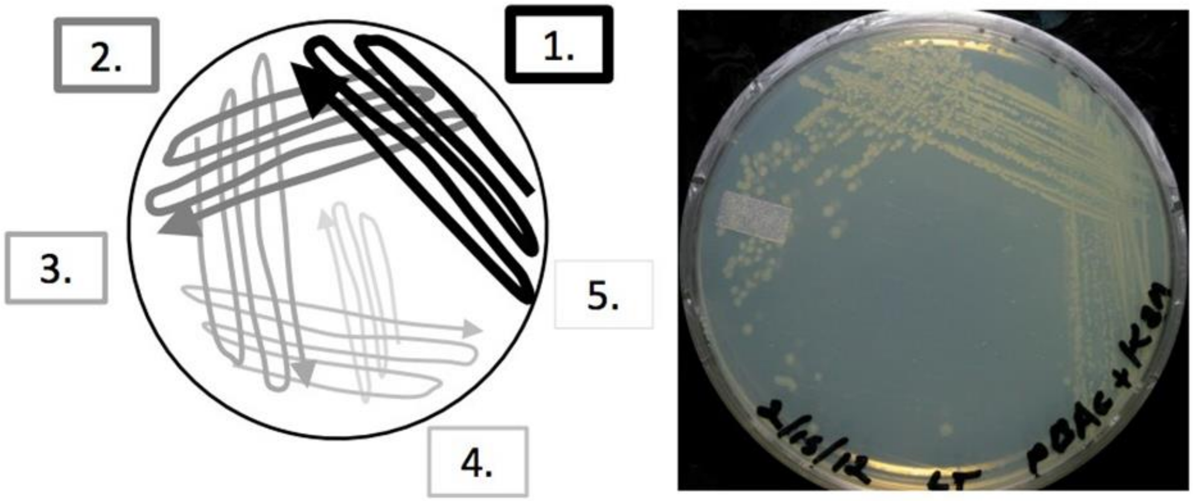

The circle with the different shade lines (image on the left), depicts pattern of spread of streaked plasmid, with image of the actual culture plate with colonies shown on the right. The number 1., 2., 3., 4. and 5. show how the streaked cells are spread from the original first tooth pick streak of DH5α cells containing the plasmid pQE_WgMDH, with the dark thick lines in pattern 1., then subsequently through to pattern 5, using new sterile tooth picks for each spreading cycle, with the declining darkness in line shades depicting decreasing amounts of cells.

#### Day 1-PP: Start/Prepare an overnight culture

##### Objectives

Today you will choose *E. coli* colonies containing the plasmid with the cloned MDH gene from streaked plates. You will use these colonies to inoculate media and start overnight cultures. Next time you will isolate plasmid DNA in a procedure called a miniprep on these cultures.

##### Procedure

DH5α *E. coli* colonies transformed with the plasmid pQE_WgMDH were streaked onto plates containing LB agar and 100 µg/mL of ampicillin. Choose large round isolated colonies for inoculation.

1. Establish an aseptic environment by spraying down your workspace and wiping with paper towels, and spray your gloves with 70% ethanol.

2. Obtain a culture tube containing 3 mL of LB. Add ampicillin to a final concentration of 100 µg/mL. The stock concentration is 100 mg/mL.

3. From a plate of transformed colonies, identify a large isolated colony.

4. Angle the lid over your plate to protect it from contamination as you reach in with a toothpick. Dab your colony with a sterile toothpick and drop the toothpick into a culture tube containing 3 mL of LB plus 100 ug/mL of ampicillin.

5. Label your culture tube with the contents (strain, vector), your class and section, your group number, and your initials. Incubate overnight at 37° C shaking at 190 rpm. The next lab section you will return to these cultures to observe whether there was growth and perform a plasmid miniprep.

### Day 2-PP: Plot growth curve of *E. coli*

#### Objectives

Be sure to start the growth curve at the beginning of lab to ensure you have sufficient time to complete it. The objective is to observe the pattern of E. *coli* growth. You will produce a growth curve by plotting the absorbance, sometimes called optical density (OD), measured at 595 nm against time, of a growing bacterial culture (In this case, *E. coli* DH5α). You will then be asked to identify the different phases of growth and determine the optimal period in which one would add the transcription inducer, isopropyl β-D-1-thiogalactopyranoside (IPTG).

#### Background

Recall that the objective of Boot Camp is to express the gene for MDH, purify MDH in large amounts, and characterize its enzymatic function. We will be using E. *coli* BL21 (note that DH5α is used for plotting the growth curve, but BL21 which is the expression plasmid/vector, is used for expressing the gene into the MDH protein later), as the host expression cell line. IPTG is an analog of lactose, and because our gene of interest is under the control of the lac promoter, addition of IPTG will induce expression of our gene of interest.

In order to obtain large amounts of MDH, it is critical to induce expression of the MDH gene when growth of the E. *coli* is optimal. As a newly diluted culture of E. *coli* begins to grow, it starts out slowly, but as long as it has enough nutrients, it will generate the proteins needed for cell growth, and reach a maximal growth rate at which the number of cells is increasing exponentially with time. This is known as the exponential or log phase of growth. Induction is generally performed at mid-log phase, or when the culture is well into logarithmic growth, but before it begins to use up all the nutrients in the medium. Once induced, plasmid-containing E. *coli* cultures often show slower growth as they direct most of their resources to making the expressed recombinant protein at the cost of growing rapidly.

Figure 3: Bacterial growth curve over time, showing the different phases of growth.
Source: Bertrand, J. Bacteriol. 2019; doi:10.1128/JB.00697-18

**A repres*entativ*e growth plot of a bacterial culture.**

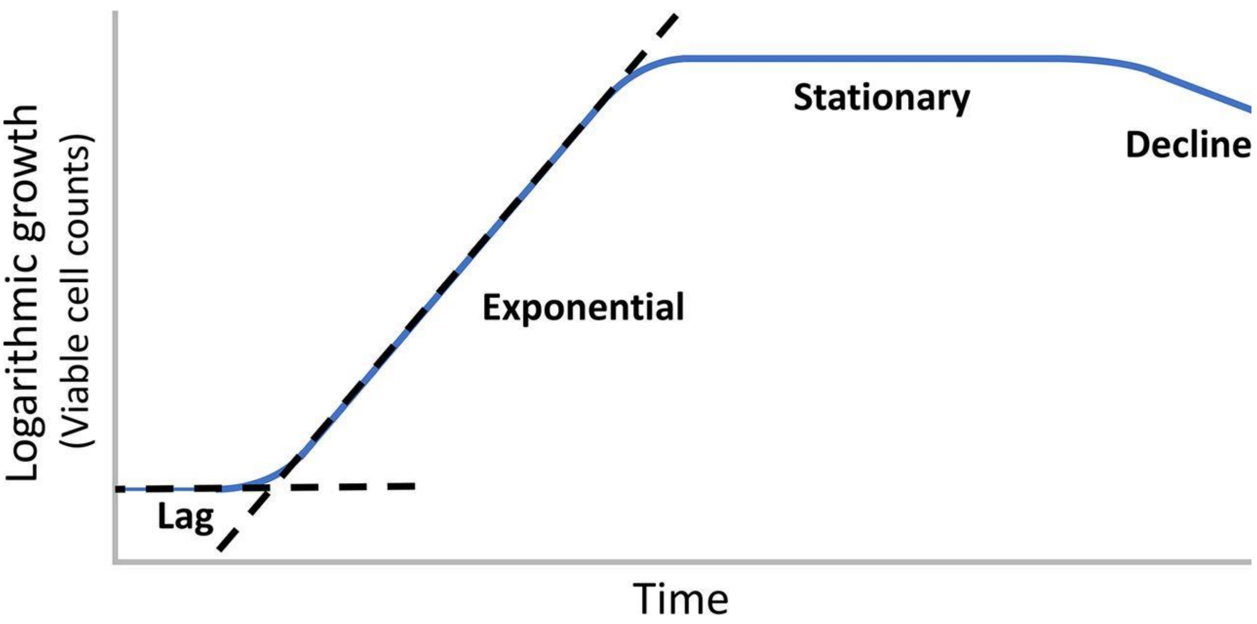

You will monitor growth over time by measuring the absorbance of each culture at 595 nM, using a spectrophotometer. This apparatus measures the amount of light of a given wavelength that passes through a sample. The less light that passes through, the greater the absorbance. You will plot your results and make note of the: lag phase, when the culture is initially growing slowly, the log phase which is exponential, the transition to stationary phase as nutrients are depleted and eventually decline phase, as cells die due to lack of nutrients.

Cuvettes: Absorbance (optical density) is read from your samples in an optically clear cuvette. Because light will be passing through the cuvette, any scratches, smudges, or fingerprints that you leave on the clear side of the cuvette can scatter light and affect the accuracy of your measurement. Be sure to know which sides of the cuvette need to be kept clean and handle these cuvettes carefully.

#### Procedure

1. Obtain a flask with 25 mL of LB. Label your flask with the contents, (the strain of E. *coli* is DH5 α), your class and section, your group number, and your initials.
2. Obtain a culture tube of grown *E. coli strain* DH5α from the shaker. Inoculate your LB-containing flask fresh LB with 0.5 mL of this *E. coli* culture.
3. Take a baseline reading of the 25 mL culture you just inoculated. When you obtain two 1.5 mL visible light cuvettes, do not touch the clear/transparent side where light will pass through. Add 0.5 mL of inoculated culture and 0.5 mL of LB to 1 cuvette and 1 mL of LB to a second. Measure the absorbance of the culture against the LB as a blank. Record the time and the absorbance and plot this point. Remember to multiply your reading by two to account for the ½ (1in2) dilution. If your baseline reading absorbance is below 0.05 at 595nm ask an instructor.
4. Put your flask on the shaker at 37° C and at least 190 rpm. This can be done as soon as 0.5 mL of culture is removed for the absorbance measurement. Minimizing the time your culture is at room temperature will improve your growth curve.
5. Fill the cuvette that contained culture with 70% ethanol and let it sit for ten minutes. Then rinse your cuvette with reverse-osmosis deionized water from the white tap in the middle sink until you can see no remaining LB.
6. Repeat your measurements regularly, every 30-45 minutes. As the rate of growth increases, take measurements more frequently. Minimize the time your cultures are left at room temperature by leaving your flasks on the shaker as you remove your sample. When you have finished, discard your LB in liquid waste and rinse cuvettes with reverse-osmosis deionized water.
7. At the end of lab, gently place your clean cuvettes in the Styrofoam box or on the peg rack on the bottom of the cart to dry. Show your graph to an instructor and identify each stage of growth and the ideal induction range.

### Day 2-MB: Plasmid DNA preparation (Miniprep) of inoculated cultures and restriction digestion

#### Objectives

The first objective is for each group member to purify plasmid DNA from your respective cultures inoculated on Day 1. The second objective is to perform restriction digestion on these samples to verify that they contain the correct plasmid.

#### Background

##### Plasmid Purification

Each bacterial cell has one main, circular chromosome that carries most of the bacterium's genes. In addition, a bacterial cell may also contain a much smaller, circular DNA molecule called a plasmid. Plasmids are copied separately from the main chromosome and can accumulate to levels of a few to many copies per cell. In nature, plasmids may carry one or more genes, such as antibiotic resistance genes.

We will be using a technique called alkaline lysis to purify plasmid. Cells are concentrated through centrifugation and re-suspended in Solution 1 (50 mM glucose, 10 mM EDTA, 25 mM Tris-HCl at pH 8.0). Glucose acts as an osmo-protectant, preventing premature cell lysis. The EDTA (ethylenediaminetetraacetic acid) stabilizes the DNA backbone and reduces DNAse activity by chelating magnesium ions required for the activity of most nucleases. You will also add RNase A, a pancreatic ribonuclease that does not require a metal cofactor and therefore remains active in solutions containing EDTA. Cells are lysed in an alkaline solution 2, (1% sodium dodecyl sulfate and 0.2 M NaOH). Sodium dodecyl sulfate (SDS) is an ionic detergent that disrupts cell membranes. The high pH of the 0.2 M NaOH denatures macromolecules. The reaction is neutralized with Solution 3 (3.0 M potassium acetate, pH 5.5). During this step, plasmids renature successfully while proteins and chromosomal DNA do not and are separated by centrifugation. Isopropanol and ethanol are used to remove salt and pellet plasmid out of solution. Plasmid is often stored in TE buffer (10 mM Tris-HCl, 1 mM EDTA, pH 8.0). Different strains of *E. coli* are optimized for different research applications. DH5α is a strain of *E. coli* optimized for long-term cryogenic storage and for the production of many copies of a plasmid. As such, you will perform a miniprep DNA extraction with the DH5α strain of *E. coli* transformed with pQE_WgMDH.

##### Restriction enzyme digestion

The full-length MDH cDNA is ~1000bp and the restriction sites added to the PCR primers are NcoI and Bgl II. For sub cloning to be successful, the same restriction enzymes must be used to digest the expression plasmid pQE60. Restriction endonucleases cleave double stranded DNA at a site in very specific palindromic sequences (usually 4, 6, or 8bp recognition sequences). These enzymes are produced naturally by bacteria (likely as a defense mechanism to digest foreign DNA), and play a critical role in molecular cloning. As the sequence recognized by a restriction enzyme is cut in a specific way, and because DNA structure and composition is essentially universal, DNA from one source (such as a human gene) can be sub cloned into another source (like a bacterial expression plasmid). The restriction enzyme generates compatible ends in each fragment; these ends can specifically bind to each other and join the two DNA fragments through base pairing. DNA ligase is an enzyme that covalently links the DNA phosphate backbones of the two molecules, and is necessary for permanent insertion of target DNA into plasmids.

#### Procedure

##### Miniprep of Plasmid

1. Obtain your 3 mL cultures of *Escherichia coli* (E. *coli)* transformed with the plasmid pQE-WgMDH.
2. Transfer cultures to separate micro centrifuge tubes. Spin the tubes at full speed for 30 seconds. Verify the cells fully pelleted the discard the supernatant.
3. Repeat Steps 1-2 for the remainder of the 3 mL cultures, or as much as you are able to retrieve from the culture tube.
4. Vortex the tube containing the cell pellet to re-suspend the pellet in the small amount of liquid remaining.
5. Add 100 µL of Solution 1 (50 mM glucose, 10 mM EDTA, 25 mM Tris-HCl at pH 8.0), and vortex to mix.
6. Add 10 µL of 1 mg/mL Ribonuclease A (RNase A); pipet gently up and down using the same pipet tip you used to add the RNase. To mix thoroughly, cap the tube, and turn it upside down several times.
7. Add 200 µL of Solution 2 (1% sodium dodecyl sulfate and 0.2 M NaOH) to each tube. Close the cap, and mix the solution by rapidly inverting the tube a few times. Do not vortex, because at this stage vortexing would shear the chromosomal DNA into smaller fragments, which would then contaminate your plasmid sample.
8. Let the tube stand on ice for 5 minutes. It is fine if this step goes longer.
9. Add 150 µL of ice-cold Solution 3 (3.0 M potassium acetate, pH 5.5) to each tube. Close the caps, and mix the tubes by quickly inverting them a few times. DO NOT VORTEX. A white precipitate will form.
10. Let the tube stand on ice for 5 minutes. It is fine if this step goes longer.
11. Place your tubes into a micro centrifuge so that they are balanced, and spin the tubes at full speed for 5 minutes. The precipitate will pellet along the side of each tube.
12. Carefully transfer the supernatant (top, clear liquid, which contains the plasmid) into a clean, labeled 1.5 mL tube, being careful not to pick up any of the precipitate. KEEP the tube with the supernatant, and discard the tube with the precipitate.
13. Add an equal volume (about 400 µL) of 100% isopropanol to precipitate the plasmid DNA.
14. Immediately close the cap, and mix thoroughly by inverting the tube several times.
15. Let the tube stand on ice for 2 minutes.
16. Then, with another group, place your tubes, with their hinges pointing outward from the center, in a micro centrifuge (balanced), and spin at full speed for 5 minutes.

17. Carefully remove and discard the supernatant (liquid). The reason that you oriented the tubes with the hinges outward is that the plasmid DNA pellet may be difficult to see. It will be at the bottom, on the hinge side of the tube.
18. Add 200 µL of 100% ethanol to each tube, and mix by inversion several times.
19. Spin your tubes at full speed in a micro centrifuge for 2 minutes (hinges out).
20. Carefully remove and discard the supernatants. Use a P20 or P200 pipettor to remove as much liquid as possible without dislodging the pellet of plasmid DNA.
21. Re-centrifuge the tubes for about 10 seconds, and remove the liquid with a pipet.
22. Allow the remaining traces of ethanol to evaporate for 5-10 minutes.
23. When the ethanol is gone, add 20 µL of TE buffer (10 mM Tris-HCl, 1 mM EDTA, pH 8.0) to each tube to re-suspend the DNA. Gently pipet the solution up and down a few times over the pellet and the tube wall under the hinge, to ensure that all of the plasmid DNA comes into contact with the TE buffer.
24. Centrifuge the tubes briefly (5-10 seconds) to collect the liquid at the bottom.

25. Make sure that each tube is labeled with the name of the plasmid, your group number, today's date, and your initials.
26. Place the labeled tubes of plasmids on ice, for temporary storage until you are ready to set up the restriction digestion reactions.

#### Restriction Digest

1. Choice of restriction enzymes. Choose 2 enzymes that will give a predictable digest pattern (hint, do not just use the two enzyme that were used for cloning- what other sites are there? use the map to make your choices.)

**List of suggested enzymes available**

| | |
|---|---|
| Apa I | NdeI |
| Bgl II | Xba I |
| EcoRI-HF | Xho I |
| Hind III-HF | XmnI |

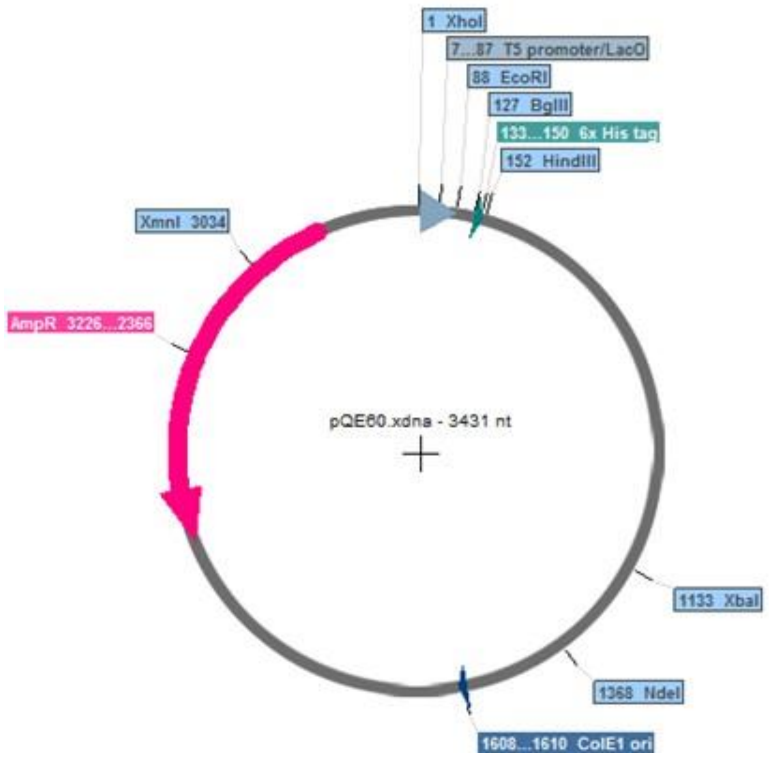

Figure 4: Map of pQE60 showing restriction sites (made using Serial Cloner)

2. Set up your digests with a final reaction volume of 20 µL. Each tube will contain DNA (from your miniprep), restriction enzyme(s), buffer and water to a final reaction volume of 20 µL. Each reaction must contain 1/10th volume of the 10X reaction buffer and 1 µL of each enzyme solution. Note that the final glycerol concentration cannot exceed 5% (enzymes are stored in 50% glycerol). Since you do not know the concentration of your plasmid DNA, we provide an estimate of what amount to use based on previous students' experience in the lab. You should use 2-4 µL of DNA sample per reaction. Enter the volumes for each component in the table below, and indicate which DNA and which enzyme for each. (Add more columns if you have more than two reactions).

3. Mix both tubes by flicking with your finger and incubate at 37°C; teaching staff will transfer to -20°C after at least one hour.

4. Mix both tubes by flicking with your finger and incubate at 37°C; teaching staff will transfer to -20°C after at least one hour.

5. For future experiments, you can vary how these digests are set up, you can use 10 µL reactions to save reagents and DNA, and if you are setting up several reactions with the same enzymes, you can make a master mix of all the reagents, then aliquot that to add the different DNA samples. Also, for high quality DNA samples with no RNA contamination, you can use lower ng amounts of DNA per sample.

#### Table 2: Restriction digest reaction components

| **Component** | **Reaction 1** | **Reaction 2** |
|---|---|---|
| DNA | µL | µL |
| 10X buffer | µL | µL |
| Enzyme 1 | µL | µL |
| Enzyme 2 | µL | µL |
| dH2O | µL | µL |
| **Total volume** | **20 µL** | **20 µL** |

### Day 3-PP: Large-scale induction

#### Objectives

Today you will perform a large-scale induction. You will induce expression of MDH by adding IPTG to a culture of BL21 E. *coli* that contains pQE_WgMDH.  Induction is performed after the culture has reached log-phase growth. Begin the induction procedure at the beginning of the lab period.

#### Background

For many research applications, it is advantageous or even necessary to have a large supply of the purified protein under study. In the past, proteins were purified exclusively from their natural host organisms. To obtain large amounts of the protein, a researcher had to have large amounts of the host organism at their disposal. Even in cases where that is readily available, it is difficult to purify sufficient amounts of protein encoded by genes that are expressed at low levels. In the case of purifying human proteins, it is difficult to obtain sufficient amounts of host organism at your disposal.

It was a great step forward when cloning and transformation procedures were developed that permitted genes from any organism to be introduced into bacteria like *E. coli*. To express a foreign gene in *E. coli,* it is necessary to provide the gene with the transcription signals recognized by the bacterial RNA polymerase, since these "promoter elements" differ in structure from those used in eukaryotes.  When the first foreign genes were ligated to *E.coli promoter* elements and transformed into *E. coli,* however, the *E. coli* cells did not grow.  Since the cells were able to grow when transformed with a vector lacking the coding sequences of the foreign gene, researchers concluded that many foreign gene products are toxic for the E. *coli* cell.

However, several genes in *E. coli* are only turned on or induced under certain growth conditions. Researchers reasoned that if the foreign genes were ligated to promoter elements obtained from these inducible genes, the *E. coli* host cells should be grown to a high density before turning on the expression of the potentially toxic foreign gene for a short period of time. The cells could then be harvested before the protein had a chance to exert its toxic effects. This approach was successful, and for many years, foreign genes have been expressed in *E. coli* under the control of various inducible  *E. coli* promoters, such as lac, tac, and pL.

1. The BL21 strain of *E. coli* is optimized for protein expression. Our specific construct, the pQE-60 vector from Qiagen, uses the lac operator to control expression. Induction of expression is achieved by adding an analog to lactose, IPTG, to the cell culture. Without IPTG present, the lac promoter has a repressor bound, blocking transcription of the genes it controls. When IPTG is present, the repressor will detach from the promoter, allowing the genes following to be transcribed. Induction is ideally performed when the culture has an absorbance in the range of 0.5-1.0.

##### Procedure

2. Obtain a 250 mL flask containing 55 mL of LB. Label your flask with the contents, (the strain of *E. coli* is BL21), your class and section, your group number, and your initials. Add ampicillin to a final concentration of 100 µg/mL. The stock concentration is 100 mg/mL.
3. Transfer 1 mL of the BL21 *E. coli* pQE_WgMDH overnight culture to 55 mL of LB plus 100 ug/mL of ampicillin.
4. Begin with a baseline reading as described for the growth curve in Day 2. (If your baseline reading is below absorbance of 0.1 at 595nm, you may need to add more of the overnight culture. Otherwise, continue to the next step.)
5. Monitor the culture as you did for the growth curve. Your objective this time is to determine when it reaches mid-log growth phase (absorbance of ~0.5 at 595 nm). This should take about 2 hours at 37° C shaking at 190 rpm. Remember to multiply your OD values by the dilution factor.
6. At the point just before induction, take 1 mL of culture in a micro centrifuge tube and spin at maximum speed for 5 min in a micro centrifuge, discard the supernatant liquid, and save the pellet at -20° C, important control for later analysis.
7. Induce your culture with IPTG at a final concentration of 1 mM.
8. Now move your flask to the room temperature shaker, and incubate your induction overnight at room temperature, shaking at 190 rpm.
9. In the morning, the TAs will spin down the cells from the culture and freeze them for you to use for the purification.

#### Day 3-MB: Transformation of bacterial cells

##### Objectives

The objective is to transform DNA into the DH5α strain of *E. coli* cells, treated to be competent. The plasmid used in the transformation is pQE_WgMDH, used at two different final concentrations. You will also transform a control plasmid, pUC19. After your transformed cells have been plated and allowed to grow overnight in the incubator, you will be able to count the number of colonies and determine the transformation efficiency.

##### Background

A transformation is the process of inserting DNA into a particular strain of bacteria. These bacterial cells have been treated to more readily take of DNA in their environment and are thus termed “competent” cells. To confer competency, bacterial cells arrested in the log phase of growth are chilled and incubated in a high salt environment, making the cell membranes more porous to DNA, then frozen at -80°C. Transformation with pUC19 plated on a plate with an antibiotic in this experiment is a control for the integrity of the DNA. Plating this control transformation onto an additional plate without antibiotic controls for the viability of the competent cells.

#### Procedure

You will be given 4 tubes containing 100uL of competent bacterial cells on ice. You will be transforming, into separate tubes: 2 samples of pQE_WgMDH DNA, one at 20 pg, one at 2 ng, and one sample of 20 pg supercoiled pUC19 as a control. **All tubes of bacteria must remain on ice! *If the cells remain at room temperature for even 1 minute, viability will be compromised!***

1. Thaw tubes of competent cells on ice without letting them warm up. GENTLY flick the sides of the tubes to see if they are liquid, and then quickly return them to ice.

Add each DNA sample to a separate tube of competent cells:
20 pg of pQE_WgMDH
2 ng of pQE_WgMDH
20 pg of pUC19 The final tube is a control and does not need DNA.
*Note, or you can use 3 tubes, and spread pUC19-transformed cells onto the null plate for viability.

2. Incubate tubes on ice for 10 minutes.
3. Carry the tubes (in the ice bucket) to the water bath. Incubate at 42°C in water bath for exactly 45 seconds to heat shock the cells.
   ***Note! move directly from ice to 42°C!***
4. Incubate on ice for 1-2 min.

Working carefully to minimize contamination, add 300uL of SOC (Super Optimal Broth) media to each tube of cells.

Place reaction tubes in the 37°C shaking incubator for 60 minutes. Tubes should be closed firmly and taped horizontally to the shaking platform.

Label 4 LB + Amp plates on the bottom around the edge - do not label lids as they can easily be switched! The pUC19 control will be plated on a LB + Amp plate as a positive control. Labels should contain your course number and section, group number, date, strain and vector, and the volume plated.

Using aseptic technique, plate ~200 µL volume of each transformation solution on the corresponding plate(s). Ask an instructor if you need a demonstration. Spread cells evenly and thoroughly over the plate. Use a new sterile spreader for each plate!

Incubate plates in 37°C incubator overnight. **Note: place in incubator LB-side down for 10 minutes to allow cells to sink into agar, and then flip upside-down for the remainder of the incubation.** Plates will be moved to 4°C storage tomorrow by teaching staff.

### Day 3-MB: Agarose gel for restriction digests

Today you will perform agarose gel electrophoresis to analyze the restriction digestion you performed on your plasmid DNA.

#### Background

Gel electrophoresis is used to separate DNA so that their sizes can be measured. DNA has a net negative charge due to the phosphate groups along the backbone. In an electrical field, DNA will migrate toward the positive pole. Agarose is a gelatinous medium that restricts the movement of large DNA molecules more than small DNA molecules. Including a set of DNA fragments of known size (markers) in one lane allows the sizes of unknown DNA fragments in other lanes to be easily estimated.

#### Procedure

**Step One: Prepare the Agarose Gel**

1. Using a 250 mL flask, prepare a 1% w/v agarose gel in 50 mL 1X TAE. Weigh out ______g agarose and add to 50 mL 1x TAE. Note: two groups can share one gel.

2. Melt in microwave until no visible grains remain (use oven mitts to hold the flask) (approximately a minute, but ensure it does not boil over).

3. Allow the agarose to cool to ~60°C (warm but not burning to the touch with your gloves on).

4. SYBR Safe is at a 10,000X concentration. Add ____µL SYBR Safe dye per 50 mL of gel and mix by swirling. Pour gel solution as demonstrated by TAs.

5. Do not disturb gel until it has solidified.

**Step Two: Prepare and Load DNA Samples**

**Note: How much of your DNA sample to add is determined by the type of DNA sample you are analyzing. In this case, you have only a small amount of DNA in your digestion reactions, so you will load all of the DNA from that tube onto the gel. Sometimes you have highly concentrated DNA in your sample, and then you would load only a fraction of it onto the gel.**

1. To a new tube, 1 uL of your prepped plasmid, and the appropriate volume of 6X loading dye to a final concentration of 1x.

2. Prepare Digestion reactions to be analyzed by adding the appropriate volume of 6X loading dye to a final concentration of 1x for each digestion tube (this will increase the volume in the tube). The volume of your digest reaction is ______. Add ______ of 6X dye. Note that

you will not dilute your digests with water because there is a lower concentration of DNA in these tubes than in for example a miniprep or PCR reaction.

3. When the gel has solidified, remove the comb, and orient the gel according to the flow of the current. Submerge the gel in 1X TAE buffer.

4. Into 1 well, load your molecular weight marker. (If your expected product size is less than 1 Kb, use the 100 bp molecular weight marker, if it is greater than 1 Kb, use the 1 Kb ladder).
The concentration of DNA in each of these markers is 50ng/µL. You should load 250 ng DNA.

5. Load at least 15 µL of digestion reaction per well. Load all of your plasmid with dye sample to another well.

**Step Three: Run Gel**

Place the cover on the gel box, hook up the leads, and set the power supply to 150 volts. Check to ensure you see bubbles rising from the thin wires in the electrophoresis tank to confirm that the current is running properly. Cover your gel with aluminum foil to protect SYBR Safe from light.

1. Switch the gel off when bromophenol blue dye has migrated one-third to halfway down the gel.

2. After electrophoresis, handle gel with gloves! Photograph the gel under UV light.

3. Analyze your gel: Label each lane and write down the sizes of bands you observe in each lane. This will allow you to determine 1) if your miniprep digestion reactions worked and 2) confirm that the pQE60 contains the MDH gene.

The Molecular weight marker sets (ladders) we are using are from NEB.com, and the sizes are posted on the fridges in lab. Be sure to familiarize yourselves with the sizes when you analyze your gel.

#### Gel interpretation

Interpreting gels can be challenging, especially if some of the digests were "partial" or not complete. In each tube, there were many molecules of identical DNA and many molecules of enzyme. If too much DNA is added, then the enzyme may lose activity before it finishes digesting all of the DNA. Alternatively, if not enough active enzyme molecules are added, some of the DNA will remain undigested. If this happens, some additional bands will be visible. If the DNA is being cut at a single site, then a partial digest will include some uncut DNA. If the DNA is being cut at two sites, then a partial digest may include some single-cut DNA and some uncut DNA. In order to identify these additional bands, if they are present, you will use bands in other lanes that migrate at the same rate. This is why you included a control lane of uncut DNA.

In *E. coli* cells, plasmid DNA exists as supercoiled molecules. Supercoiling allows the cells to accumulate many copies of the plasmid in a very small space. When using the alkaline lysis miniprep procedure to isolate plasmids, the high pH at one-step often nicks some of the DNA. A nick is a break of a phosphodiester bond in one but not both of the strands of DNA. Nicked plasmids are still circular, but free rotation around the remaining intact strand results in a loss of supercoiling; the resulting plasmids are termed “relaxed circles”. Therefore, your plasmid minipreps probably contain at least two populations of plasmids: supercoiled circles and relaxed circles.

Interestingly, despite having the same mass, these two types of molecules migrate in an agarose gel at different rates. The supercoiled circles, being more compact, migrate faster than relaxed circles, and as a result appear to run at a smaller size than they actually are (often appearing almost half the actual size). Relaxed circles, by contrast, get entangled in the gel matrix. It is important to note that the same plasmid can run with three different sizes depending on whether it is 1) linearized by a single cut of both strands of DNA, 2) supercoiled or 3) nicked in one DNA strand (relaxed). In summary, the speed with which circular DNA molecules migrate depends on both the size and the shape of the molecule. Examples of bands corresponding to linear (L), supercoiled circular (SC), and relaxed circular (RC) plasmids are shown in the figure to the right.

Figure 5: Plasmid DNA bands on agarose gel

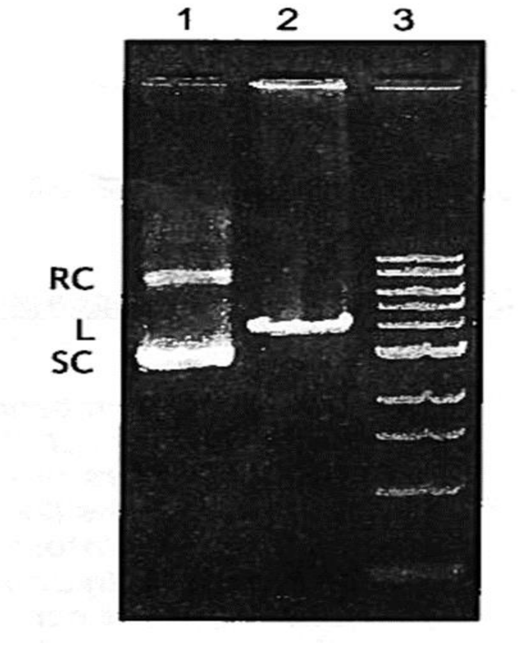

Image shows relaxed circular (RC) in lane 1, Linear (L) in lane 2 and Supercoiled (SC) plasmid in lane 1, separated into different bands. Lane 3 shows the bands from the DNA ladder.

### Day 4-PP: Cell Lysis and MDH purification

#### Objectives

The objective is to perform affinity chromatography (AC) to purify MDH away from other cellular components and proteins. We will assess the success (or failure) of the purification method quantitatively based on enzyme activity.

#### Background

##### Cell disruption to obtain cleared lysate

The induction was designed to produce as much intracellular MDH as possible. The *E. coli* must be separated from the LB through centrifugation. This will produce a cell pellet. The media must be removed and the cell pellets must then be lysed to release the MDH for further purification. There are numerous methods that can accomplish this task, each dependent on cell type (e.g., is there a cell wall present?), economics and scale. Because chemical lysis methods can be detrimental to product stability, often-mechanical methods are used, particularly at the large scale due to their cost effectiveness. You will lyse your cell pellet chemically. The addition of protease inhibitors to the lysis process can be effective at minimizing protein degradation and hence increasing yields. Benzonase nuclease or other DNases and RNases can be added to reduce viscosity of the lysed cells and increase yield.

##### Nickel Affinity Chromatography

Affinity chromatography (AC) is used to purify protein with specific affinity for the column matrix. A ligand (in this case nickel) is covalently bound to a solid support bead (typically agarose or sepharose). Recall that MDH was expressed with a his (6X-histidine) tag. The His-tag has a very high affinity for nickel, while other cellular proteins/enzymes have a low affinity for nickel. Thus, His-tagged MDH will bind specifically and tightly to the matrix while other cellular components and proteins will flow through. The bound His-fusion protein dissociates from the nickel beads by adding excess imidazole. An imidazole ring is present at the end of the histidine side chain, which makes imidazole a good competitor for binding nickel beads.

#### Procedure

Begin with step 1 of cell lysis. There are many long wait periods during the lyse step. Begin lab by lysing your cells. During the wait periods, setup your column for purification, and set up to count colonies and streak agar plates.

#### Harvesting cells (Done by Teaching Staff, and pellets stored at -80$^{o}$C until use)

1. For protein purification, you need to collect cell pellets from your MDH induction experiment. These cells should have expressed MDH at a very high level upon induction. Pour your induced culture into one 50 mL falcon tube and label with your group number.
2. The pellets will be centrifuged together with the other groups (5000 rpm for 10 min. at 4$^{o}$C in large rotor).
3. After centrifugation, the medium (supernatant) is removed, and the pellets stored until needed for lysis (pellets retrieved from -80$^{o}$C and kept on ice by TAs immediately prior to use).

#### Cell lysis

Since the MDH-His protein is found intracellularly, we must efficiently lyse the cells to recover our protein. You will also need to retrieve the 1 mL of cells you harvested in a previous lab session (pre-induction), and lyse those as well (scale the volumes of respective solutions appropriately).

1. Obtain your pelleted cells, and weigh to determine the wet mass of your cells (be sure to use an empty tube to tare/zero the balance!). Record the weight.

Wet cell mass: _______________

2. Add 5 mL of Bugbuster/gram of cell pellet. Add 1µL of DNAse for each mL of BugBuster. Add 1µL of protease inhibitor per 20 mg of cells. Re-suspend the cells immediately by gently pipetting up and down. Do not forget the 1ml of un-induced cells that you saved in the fridge during a previous lab class. You will use a similar lysis procedure, scaled appropriately.

3. Incubate at room temperature for 15 minutes on a slow shaking platform.

4. Centrifuge at 17,000 X g for 10 minutes.

5. Transfer 600 µL of the supernatant from your induced cells to a new tube and save/store at $4^{o}$C for activity assay, Bradford assay, and SDS-PAGE. **Saving your cleared lysate is CRITICAL for assessing your purification!**

Transfer the remaining supernatant from your induced cells to new tubes and save these on ice until ready for the purification. Record the total volume of supernatant at this step _______________mL

6. For your uninduced sample, transfer the supernatant to a new tube and save/store at 4° C, to be used later for your SDS-PAGE gel.

##### Calculating Resin Volume for Affinity Chromatography (AC)

You will be given 4 mL of suspended resin. This will result in about 2 mL of compact resin. You will calculate whether this is a sufficient amount of resin for your purification.

Given the original culture volume, we are purifying from, and that previous studies show that a 10 mL culture yielded 1-2 mg MDH; we can roughly estimate how many mg of MDH our purification should be able to yield. The nickel resin we are using has a stated binding capacity of 5-10 mg protein per mL resin; therefore, you can determine how many mL resin are needed to bind all the MDH. We choose an amount that will balance the need to have plenty of capacity so we don't lose any protein of interest, with the need to minimize non-specific protein binding that can occur when significantly more resin than necessary is used (and also, to minimize costs, since the resin is pricey).

Calculation: Volume of resin to bind ideal yield of protein: __________________ mL

There are two approaches for binding target protein to the affinity resin: in batch mode or through on-column binding. In batch, the pre-equilibrated resin is mixed with the impure protein fraction in a tube to allow binding. A different approach is to load the pre-equilibrated resin into the column, then apply to and pass over the impure protein to the resin. On-column binding is a standard approach, but historically batch binding has been shown to be superior to on-column binding. During Boot Camp, we will be performing on-column binding. **Begin the following protocol at Step 5 of On-column Binding.**

#### Affinity purification protocol

*During Boot Camp, begin at Step 5 of On-column Binding.*

##### Preparing the column

1. You will be provided with an empty polypropylene column, as well as a 5 mL pipet.

2. Cap the bottom of the column and drop column into a 15 mL tube.

3. Fill the column with RODI water, float the filter disc, and force the filter disc to the bottom of the column using the back end of a cell spreader or 5 mL pipet.

4. Inspect the filter in the column- it should be level and there should not be air bubbles below or above disc. Reposition as needed. If air bubbles persist, a surfactant can be added.

##### On-column binding

1. Obtain 4 mL of fully re-suspended resin solution in column buffer.

2. Open your prepared column (see above), and discard the water.

3. Pour resin into column (make sure the stopcock is closed!) using a plastic Pasteur pipet. Open the stopcock and allow liquid to flow out of the column, thereby packing the resin. Do not save this effluent.

4. Wash column with 10 resin volumes of elution buffer (NPI 250) then 20 resin volumes of His-binding buffer (NPI 10). Collect in your liquid waste. (A "resin volume" is the volume of your settled resin; recall we used ~2 mL of settled resin.)

5. Place a tube labeled flow-through under the column; this tube will be used to collect your flow-through fraction (SAVE ON ICE!). Apply the cleared lysate to column, and open the stopcock to allow a flow rate of not greater than 2.5 mL/minute. If your rate is too fast or slow, tell a lab instructor to check your column.

6. Proceed to step 7 of "Batch binding" protocol below.

**Batch binding**

1. Combine the resin and the cleared lysate in a 15 mL tube and mix gently.

2. Mix for 30 min. at 4ºC/25ºC to allow the affinity-tagged MDH to bind to the resin.

3. Open your prepared column (see above), and discard the water.

4. Place the end cap back on the column, and load your resin into the column using a plastic Pasteur pipet.

5. Place a 15 mL tube labeled flow-through under the column and collect your flow-through fraction (SAVE ON ICE!).

6. Calculate your column flow rate by adding a known volume and timing how long it takes for this volume to get to the top of the resin (but do not let top of the resin dry off). An ideal flow rate is 2.5 mL/ min.

7. Record the Flow-Through volume in your 15 mL Tube: ________________. Remove 1 mL of Flow-Through to a new tube and save/store at 4ºC for later use (activity assay, Bradford assay, and SDS-PAGE).

8. Wash the column with 20 resin volumes (Recall ~2 mL of settled resin was used.) of wash buffer (NPI 20), and collect liquid in a 50 mL tube. Record the volume of wash you used. Save/store 1mL of wash from the *end of the wash process*, store at 4ºC for later use.

9. Elute the affinity-tagged MDH from the column with 4 x 5 mL (i.e. add 5mL NPI 250, to column and collect eluted liquid in 15 mL tube, then repeat 3 more times to get 4 elutions in 4 separate tubes) of His-elution buffer (NPI 250) (SAVE ON ICE!). Save each (1 to 4) elution in a separate 15 mL tube and store ON ICE.

10. Make sure you have saved/stored 600µL of cleared lysate,1 mL of your flow-through and wash samples and the entire volume of your elutions at 4°C. Also, save ALL of your elution volume in 15 mL tubes.

11. Your TAs will wash your resin for future use in the following sequence.
    a. 10 column volumes of wash buffer,
    b. 10 column volumes of water,
    c. 10 column volume of 20% EtOH,
    d. Collect in liquid waste.
    e. Column stored in 20% ethanol, 80% NPI-10 enough to make sure resin does not dry out

### Day 4-MB: Count colonies and streak Agar plates

#### Objectives

You will refer back to your transformation plates to select isolated colonies for liquid culture. You will inoculate liquid cultures that will be saved/stored at 4°C, and then incubated at 37°C on a shaking platform overnight before the next lab. These liquid cultures will be used for culture PCR on Day 6.

#### Protocol

Each group must select five colonies for inoculation. Follow the protocol from Day 1 of the PP-stream for directions on inoculating liquid media. Be sure to label your culture tubes clearly and thoroughly.

### Day 4- Check your transformation plate for single, isolated colonies

1. Obtain your plate (Figure 6.) from where you transformed your cells with MDH plasmid, count the individual colonies, and then record in the table below.

##### Figure 6: Transformed *E.coli* cells cultured on LB Agar + Ampicillin

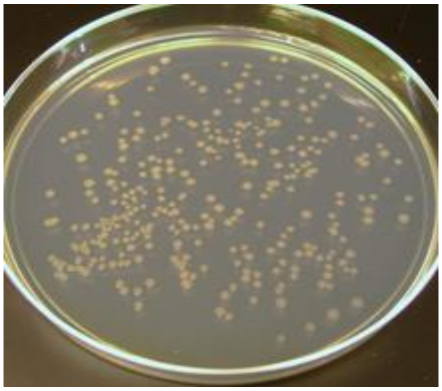

LB Agar + Ampicillin plate showing growth of single colonies on a petri dish.

Record in your notebook how many colonies you obtained for each transformation. If you have too many colonies to count the whole plate, count a fraction (1/4, 1/10, etc.) of the plate that best represents the distribution of colonies, and multiply the number obtained as appropriate.

| Transformation | # colonies obtained |
|---|---|
| 20 pg pQE_WgMDH transformation | |
| 2 ng pQE_WgMDH transformation | |

Transformation efficiency can be calculated using the positive control transformation. Efficiency is expressed as the # of colonies per µg DNA. An acceptable transformation efficiency is 1.0 x $10^7$ cells per µg DNA. Remember, a picogram is a millionth of a microgram, and 20pg of positive control DNA was utilized. After transformation with the positive control plasmid, you only plated 200 µL out of 400 µL total volume, so be sure to account for this in your calculation.

| Transformation | # colonies obtained | Transformation Efficiency |
|---|---|---|
| Positive control | | |

| Transformation | # colonies obtained | Transformation Efficiency |
|---|---|---|
| pQE_WgMDH | | |

Is the transformation efficiency for pUC19 or pQE_WgMDH greater? Why might this be?

2. Establish an aseptic environment on your lab bench.
3. With a sharpie on the bottom (agar-side), divide a new LB+AMP plate into 5 wedges, one for each colony you select for PCR. Label each wedge with a number.
4. Take a sterile toothpick and pick a single, well-isolated colony from your transformation plate with MDH plasmid: Gently dab the toothpick against the colony (No need to scrape up all of the colony).
5. Dip the toothpick lightly into a corner of the corresponding numbered section of the new LB+AMP plate. Then put another toothpick into the SAME colony, make sure you have obtained some of the cells and place this toothpick directly into the culture tube with the corresponding number. This way you can make a streak and inoculate a tube from the same colony.
6. Streak: This step is best performed after a demonstration from instructional staff. With a new sterile toothpick, go back to the corner of the LB+AMP plate where you dabbed your previous toothpick. Dip into the cells that you deposited there and gently "paint" a short streak with your toothpick (darker line in figure Figure 2 below). Take a second sterile toothpick and swipe is across the end of the streak you just made and continue further down the plate (see pattern in Figure 2 below). When you analyze your PCR results and determine which colonies contained the plasmid of interest, you will be able to access your isolated colonies from the corresponding sectioned wedge on the plate. Repeat this procedure for each colony, using sterile toothpicks each time.
7. Place your streaked plate in the 37°C incubator overnight. Be sure that your plates are LB-side up. Teaching staff will move your plate to 4°C the next morning.

#### Figure 7: Agar plate partitioned into 5 wedges for single colony cultures

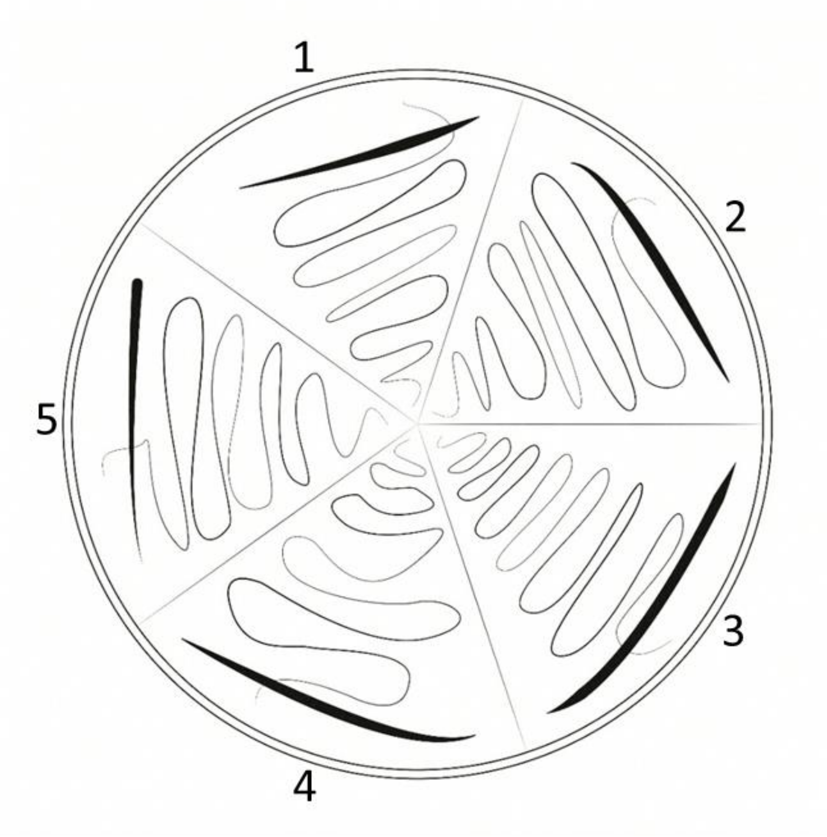

Agar plate partitioned into 5 wedges for separate single colony culture of MDH-containing transformed *E.coli* cells (as shown on Figure 4.). In each wedge the dark streak is from the first toothpick, the lighter streak is from the second too

### Day 5:-PP Enzyme activity assay for MDH activity

#### Quantifying amount of MDH

After you purify MDH, you will quantify how much MDH is in each of your purified samples collected during chromatography. We will quantify MDH using enzyme activity rather than protein level. You will use enzyme activity to determine the success of your induction and purification. Begin by calculating the total activity present in the cells; the activity found in this cleared lysate represents 100% of the total starting activity.

The ultimate goal of any purification process is to isolate the maximum amount of product, and to obtain the product in as pure a form as possible. Thus, we wish to retain as high a percentage of the starting activity as possible, while removing as much of the unwanted molecules (mainly other cellular proteins) to increase the purity of our "final' MDH preparation. Typically, total activity recovered during purification is expressed as a percent yield (recovered) based upon the starting amount of total activity. One unit (U) of enzyme activity is defined as moles of product generated per unit time (i.e., U = µmol/min.). Purity of an enzyme by convention is expressed as specific activity, the units of enzyme activity per mg of total protein. For enzymes this number reaches a theoretical maximum; i.e. if the specific activity for 100% pure MDH is 1200 U/mg protein so, 1 mg of protein that is entirely MDH will possess an activity of 1200 U. Specific activity should increase after purification (as more of the recovered protein is the desired enzyme). Note that your protein in this experiment might not have the same theoretical maximum as this example. Fold purification relates the specific activity after purification to the original (starting) specific activity, and therefore should always be greater than 1.0. Thus, any purification procedure should increase the specific activity and fold purification while maximizing the percent yield of starting activity retained.

Today you will calculate BOTH the enzyme activity AND you will use a Bradford Assay to quantify the amount of total protein (not just MDH!) isolated at each purification step. This will enable you to calculate the specific activity of MDH and the fold purification.

In the activity assay you will perform, MDH oxidizes NADH, which has an absorbance at 340 nm, to $NAD^+$, which has no absorbance, in the following reaction:
Oxaloacetate + NADH + H + ↔ Malate + NAD +
Thus, absorbance by the reactant can be used to calculate MDH activity in terms of enzyme units (moles product/unit time). However, note that you will be monitoring the activity of the enzyme by the *decrease* of NADH measured by the loss of absorbance over time. Oxaloacetate and NADH will be added in excess so as not to be limiting reagents. The reaction will take place in assay buffer (10mM K/Na Phosphate Buffer, pH 7.4). The enzyme will be inactivated by adding 1 M sodium carbonate ("Stop" solution) to stop the reaction.

#### Calculating loss of absorbance

You will set up one control with NADH and without enzyme ("NADH only"). None of the NADH in this control will be converted to NAD via MDH. You will determine the loss of absorbance due to MDH activity for each of your fractions by subtracting the absorbance for each of your fractions from the absorbance of your NADH-only control.

Absorbance of Control 2 - Absorbance of Fraction = Loss of absorbance

Use the loss of absorbance value as the $A_{340}$ value in your calculation of total activity.

##### Calculating malate dehydrogenase activity (total activity)

We are defining one unit (U) of MDH as the amount of enzyme that will convert $10^{-6}$ moles/min (or 1umol/min) of NADH at 25°C in assay buffer. You will need to convert the $A_{340}$ value into activity (umoles product formed/minute). To do this, use the Beer-Lambert equation to calculate the concentration of NADH converted (NAD+ is the product formed) per minute assay time.

The Beer-Lambert Law is represented by the equation:

$A=\varepsilon*c*l$

Where

| | | |
|---|---|---|
| A | = | absorbance (unit less) |
| ε | = | extinction coefficient ($M^{-1}cm^{-1}$) |
| l | = | path length of light through the sample (cm) |
| c | = | concentration of absorbing material in the sample (M) |

The extinction coefficient (ε) is a proportionality constant that defines the efficiency or extent of absorption and is specific for the absorbing molecule and wavelength.
The molar extinction coefficient is 6200 $M^{-1}cm^{-1}$, and the path length is 0.28 cm. Total assay volume is the volume in the micro well plate, volume of sample is calculated from the fraction of sample in the reaction, following these mathematical manipulations:

$C = \frac{A}{\varepsilon l}$ l (path length) = 0.28 cm $\varepsilon = 6200\frac{1}{M \cdot cm} = 0.0062\frac{L}{umol \cdot cm}$

$C = \frac{(A_{340})(umol)}{0.0017\ L}$ Volume in well = 0.1mL = 0.0001 L

Total umoles = $\frac{(A_{340})(umol)(0.0001\ L)}{0.0017\ L} = (A_{340})(5.8 \times 10^{-2}\ \text{umol})$

Units MDH activity=
$(A_{240})(5.8 \times 10^{-2}\ \mu mol)$/ (Assay time in min) X (Total Assay Volume in mL)/(Volume of sample in mL)

#### Protocol for MDH Assay

Notes: You will perform assays to quantify MDH enzyme activity. We suggest you perform **duplicate** analyses.

You will need to assay all fractions you obtained from AC, your cleared lysate, and your uninduced sample (determine the volume of each!). The data you are collecting is a CRITICAL component of your assessment, so make sure your assay tubes are properly set up, and your data are accurately collected and analyzed. This will allow you to generate an elution profile (U MDH activity vs. volume/fraction number) and identify the eluate fractions with significant MDH activity, which you will pool for future use.

In advance, you should prepare a data table to handle the following data you will collect for all samples - sample name; assay time (Minutes),$\varepsilon$ , $A_{340nm}$ for control 2,$A_{340nm}$ of sample; total assay volume, volume of sample MDH activity.

1. You should have samples labeled Lysate (L**)**, Flow-through (FT**)**, Wash (W**)**, Elutions 1-4 (**E1-E4** ).
2. For each elution sample you will also prepare a 1/5 dilution in **assay buffer** and assay the 1/5 dilution in addition to the undiluted samples. Use assay buffer and not water to dilute your samples.
3. Controls: these are very important. You will prepare two controls (a good idea to make each control in duplicate).

- Control 1: assay buffer is added instead of NADH (use cleared lysate as enzyme sample in control 1)

- Control 2: assay buffer is added instead of enzyme sample.

4. Combine the following components except OAA into microfuge tubes, cap and vortex. Keep your enzyme samples on ice, but incubate all other solutions at 25$^{o}$ C for 3-5 min prior to starting reaction:
350 µL assay buffer
50 µL 6 mM NADH
50 µL Enzyme sample (this is the sample you are assaying for enzyme activity)

5. Start the reaction by addition of 50 µL 20mM OAA (mix well by vortexing) * Do not add OAA until ready to start the reaction!
6. Incubate at room temperature for 2 minutes
7. Stop the reaction by adding 50µL stop solution (1 M $Na_2CO_3$) vortex and immediately place on ice to ensure reaction has stopped.
8. Load two wells per reaction: transfer 100µL of each reaction into two wells of a 96-well plate.
9. Read absorbance in plate reader at 340 nm
10. Calculate activity (below) and record in your data table. Remember you are interested in loss of absorbance and you need to calculate this first (***subtract*** sample absorbance from absorbance of the NADH-only Control).

$$\text{Units MDH Activity} = \frac{(\text{A340nm})\ (0.058\text{Mol})}{\text{Assay time in minutes}} \times \frac{\text{Total assay volume in mL}}{\text{Volume of sample in mL}}$$

10. Confer with an instructor about which eluate fractions with significant MDH activity to pool into one 15 mL tube. From this pooled fraction, you can move 1 mL into a micro centrifuge tube for easier handling during other procedures.

### Day 5:-PP Bradford Protein Concentration Assay

#### Objectives

The objective is to perform a Bradford assay to determine the amount of total protein in each of the fractions you collected during chromatography, calculate the specific activity, and fold purification.

#### Background

##### The Bradford Assay for quantification of protein concentration

The Bradford assay is a procedure based on the binding of Coomassie Brilliant Blue G-250 dye to protein in acidic solution that causes a shift in the wavelength of maximum absorption ($\lambda_{max}$) of the dye from 465 nm to 595 nm. The absorption at 595 nm is directly related to the concentration of protein in a solution, and a standard curve is generated from known concentrations of a generic protein. After addition of the dye solution to a protein sample, color development is complete in two minutes and the color remains stable for up to one hour. The Bradford method is rapid, however interference is sometimes observed with some reagents (e.g., detergents Triton X-100 and sodium dodecyl sulfate (SDS). The Bradford assay cannot determine the concentration of a specific protein in a mixture of proteins, and Coomassie binding is irreversible.

##### Creating a Standard Curve

In order to use absorbance to determine the concentration of protein for an unknown sample, a relationship between absorbance and concentration must be established. This is accomplished by creating a standard curve using linear regression. The standard curve is generated by plotting absorbances derived from known concentrations of protein. The assay system responds linearly for a limited range of protein concentrations because Bradford reagent becomes limiting and/or the range of the spectrophotometer/microplate is fixed. Because you are unsure of the concentration of the unknown, it is necessary to prepare several dilutions of the unknown sample so that you obtain at least one absorbance that falls in the linear range of the standard curve.

#### Procedure

##### Bradford Assay

You will assay your cleared lysate, flow-through, wash fraction, and elution fractions. Because the protein amounts in each fraction are likely to vary widely, you will need to assay different dilutions of each fraction so that the absorbance falls in the linear range of the standard curve. This will be accomplished using a multichannel pipettor with your unknowns. Refer back to the section on micropipetting for background on the multichannel pipettor. If you need more than one microplate, add blanks and standards to each plate!

Note: use a standard micropipettor for steps 2-4; the remaining steps can utilize the multichannel if you feel skilled enough!

Note: pipette and mix very carefully, because if you do not you may have to repeat the entire experiment.

1. Obtain a 96 well microplate and orient the plate with well A1 at the top left.
2. Add 50 µL of RODI water to wells A1 and A2. These wells will serve as a reference. The microplate reader will subtract the absorbance in these wells from all wells on the microplate (this is called blank subtraction).
3. Add 100 µL of the 500 µg/mL bovine serum albumin (BSA) standard to wells B1 and B2. **Note that stock BSA is diluted in RODI water**
4. Add 100 µL of all samples to be tested (see above) for protein concentration to wells B3-B12 as needed.
5. Pour ~ 20 mL RODI water into the tray for the multichannel.

You will need to do the next steps in two batches if you have used > eight wells in row B. Finish the first batch of dilutions, and then complete the plate by changing tips and diluting the remaining samples.

6. Add 50µL RODI water to all wells in rows C-H.
7. Locate row B. Take 50 µL out of row B wells and add to the row C wells directly underneath.
8. Mix well and take 50 µL out of row C wells and add to the row D wells directly underneath.
9. Continue this procedure until you have transferred 50 µL from row G wells into row H wells.
10. Remove 50 µL from row H wells and discard. All wells on your microplate with standard or sample now have a 50µL volume.
11. You need a volume of 160 µL in each well before adding Bradford reagent. Without touching liquid in each well, add 110 µL of RODI water to each well on the microplate, including blanks and standards. You should not need to change tips!
12. Next, add ~ 10 mL of Bradford reagent (Coomassie brilliant blue G250 dye) to reservoir and set the multichannel to 40 µL, and add Bradford reagent to all utilized wells. Careful: Bradford reagent is hazardous and stains skin and clothing!

13. Mix each well individually with a pipette until you are sure the dye reagent and sample have mixed.
14. Let the reaction stand at room temperature at least 20 minutes from adding the Bradford reagent for color reaction to stabilize.
15. Read the microplate at 595 nm and obtain absorbances (these will be blank-subtracted for you by the software) for all standard and sample wells.

#### Steps to determining the concentration of protein from an unknown

1. Set up a table to record the absorbances in each replicate well. If these duplicate reads are close in value, proceed below. If they are not, consult with the instructor. You may need to repeat you assay if your duplicates do not agree.

2. If your replicates are good, take the averages of your standard replicates and put them into a chart with the known amounts of BSA in each well. Remember that you ended up with 50 µL of 500µg/mL BSA stock solution in the first standard well.

Table 3: Protein Concentration sample data

| Amount of BSA in well (µg) | Average absorbance of replicates at 595 nm |
|---|---|
| 0 | |

3. Generate a standard curve using Excel. Select the chart you are working on. Click the Insert tab, then on the Scatter tab, and select the scatter plot style without any lines. To properly format your graph, click on the Layout tab to add titles. Use a descriptive chart title, label the axes (do not forget units!) and remove the legend. Under the Layout tab, click on Trend line to add the line of best fit. Add the equation of the line and R-squared value on the plot (this is under "more trend line options").

4. Analyze the absorbances for your unknown samples. *Make sure that the absorbances decrease as expected with increasing dilution.* Discard absorbances that do not fit this pattern. Choose values within the linear range of your standard curve, to calculate amount of protein. You have the most confidence close to the middle of the standard curve.

5. Use the equation of the line to determine the amount of protein in your well.
6. To determine protein concentration, multiply the amount of protein by the dilution factor and divide by the volume of stock sample added to the first well. For example if you selected an absorbance value from row F of your plate, you would multiply by 16 and divide by 50µL.

#### Calculating malate dehydrogenase activity (total activity)

For activity, use the values you calculated earlier under "Calculating malate dehydrogenase activity (total activity)." For percent yield: assume the activity of cleared lysate is 100%. For protein amount, use the total sample volumes you recorded on Day 4-PP: (Cell Lysis section ) when you purified MDH, to multiply by the concentration of sample you calculated today (Note that this table uses mg not µg).

#### Table 4: Data Table for Calculating Yield, Specific Activity, and Fold Purification of affinity column

| Fraction | Lysate | Flow Through | Wash | Pooled Elution |
|---|---|---|---|---|
| Activity (U) | | | | |
| Percent yield (%) | | | | |
| Protein Concentration (mg/mL) | | | | |
| Protein amount (mg) | | | | |
| SA: specific activity (U/mg) | | | | |
| Fold purification (SA fraction / SA cleared lysate) | | | | |

#### Day 5- Check your streaked plate for isolated colonies and restreak if necessary

1. Obtain your plate from last week where you streaked your colonies from the transformation plates. Did your streaks yield isolated colonies? You must have at least one isolated colony per student in your group for future experiments.

2. If you do not have isolated colonies of bacteria containing the pQE_WgMDH plasmid you will need to restreak your cells onto a new plate.

3. Establish an aseptic environment.

4. Label new LB+AMP plates for the colonies that need to be re-streaked.
Take a sterile toothpick and pick an isolated colony from your streak plate (or go back to your transformation plate if nothing grew). Gently dab the toothpick to pick up cells, streak it into a corner of the corresponding numbered section of the new LB+AMP plate (refer to figure 7, in the Day 4 instructions).

Repeat this procedure for each colony that needs restreaking, using sterile toothpicks each time.

### Day 6-MB: Culture PCR

#### Objectives

The objective is to perform PCR on cultures grown from transformed colonies to screen for the presence of your plasmid of interest. You will be using primers that flank the region containing the MDH gene.

#### Background

The components of a standard PCR reaction are the DNA template, sequence-specific primers (Two), a thermostable DNA polymerase (such as Taq), free nucleotides, and a suitable buffer with $Mg^{+2}$. The set of two PCR primers flanks the region of DNA to be amplified. Using double-stranded DNA as a template, PCR can specifically amplify the template DNA sequence exponentially ($2^n$). PCR exponentially amplifies ONLY the region of DNA between and including the two primers. While the DNA sequence information at the primer-binding site is essential to design the primers, the region between the primers does not have to be of known sequence.

PCR occurs in cycles of three sequential steps: denaturation, annealing, and extension. Denaturation uses heat (95ºC, above the $T_m$ or melting temperature of double stranded DNA) to separate the double-stranded DNA template. Lowering the temperature just below the $T_m$ of the primers (~55ºC) promotes primer annealing to the template DNA sequences complementary to each primer. The thermostable DNA polymerase catalyzes the extension of each primer using free nucleotide triphosphate (bases) directed by the nucleotide sequence of the template DNA at ~72ºC. Since PCR occurs at high temperatures that would denature conventional DNA polymerases, thermostable polymerases obtained from thermophilic organisms are utilized. The above denature-anneal-extend cycle is repeated 35-40 times, yielding millions of copies from each original template DNA molecule.

Success in PCR relies in part on the design of primers. Primer design is often complex, and primers that should work on paper sometimes fail in the PCR tube. An ideal set of primers binds only to the DNA sequence of interest (with high specificity), and do not bind to themselves or each other (primer-dimers essentially remove free primers from solution). The primers within a set can be separated by <100 bases or by several kilobases, but must be properly oriented to amplify on the antiparallel strands across the DNA region of interest. Primers in a pair should have similar $T_m$ to maximize binding specificity. It is important to note that while primers must be highly specific for the target DNA sequence, they do not need to be exactly complementary to the target sequence. Thus, PCR can be used to incorporate nucleotide changes (used in site-directed mutagenesis), or to add restriction enzyme recognition sequences to the ends of the amplified DNA sequence (as was done when MDH was cloned into pQE60). Since these same primers with the restriction enzyme site are utilized in every PCR cycle, the amplified DNA fragments will contain these engineered sites.

You will be performing culture PCR, in which a small amount of liquid culture is added to a PCR reaction as the source of template DNA. This liquid culture is grown from an independent colony (separate from, and not in contact with, other colonies on the culture plate) selected from a transformation plate, picked using a tooth pick and re-suspended directly in a PCR mix. All the cells in a colony should be identical, that is, derived from the same initial cell. During the thermocycling, the cells lyse and the plasmid DNA can serve as a template for PCR.

Recall that MDH was cloned into pQE60. Primers that either specifically recognize only the MDH insert, or that flank (surround) the MCS can be used to determine if insertion into the MCS occurred during sub cloning, simply by looking at the presence and size of the DNA fragment amplified by PCR.

#### Procedure

Each group will set up as many as 5 PCRs using your cultures grown from transformed colonies as the source of template DNA. The negative control will contain water instead of template. The positive control will contain pQE-WgMDH plasmid as a template. The primers are spaced 400 bp apart.

In total, you will be doing 7 PCR reactions: 5 on cultures + 1 negative control + 1 positive control

1. You will prepare a master mix enough for each of your PCRs plus one. Use the following table to determine how much of each component should be added

Table 5: PCR Master mix components

| **Component of 25 µL reaction** | **Concentration in Reaction** | **Volume in 25 µL reaction** | **Volume in 8x master mix** |
|---|---|---|---|
| 10x Standard Taq reaction buffer | 1x | | |
| 10mM dNTPs | 200µM | | |
| 5 µM Forward Primer TGCTTCCTGCCAACTCTTT | 0.5 µM | | |
| 5 µM Reverse Primer GATCCTCGGGATGTTGATGTT | 0.5 µM | | |
| Taq polymerase 5000 U/mL | 1.25 U/ 50 µL rxn | | |
| Autoclaved reverse-osmosis deionized water (round to nearest µL) | ------------------------- |  |  |
| Total volume of master mix | ------------------------- | 24 µL* |  |

*The volume of master mix for each 25µL reaction is 24 µL. The remaining volume per reaction is for the source of template. Template does not go into this master mix.

2. Make your 8X master mix, adding the Taq polymerase last.

3. Label your 7 PCR tubes (group ID-reaction number), and put them on ice. Add 24 µL of the 1X PCR Master Mix to each tube.

4. Add 1 µL of water to your negative control reaction.

5. Add 1 µL of each liquid culture to the corresponding experimental reaction tube.

6. Add <1000 ng of pQE-WgMDH template DNA to your positive control tube. The stock concentration is 500 ng/µL.

7. Place your tubes in a thermocycler and run with the following program. This program has been entered into the thermocycler in Room 260 under the name MDH400BP

Table 6: PCR cycle components

| # Cycles | Temperature (°C) | Duration |
|---|---|---|
| 1 | 95 | 4 min |
| 40 | 95 | 10 sec |
| | 57 | 30 sec |
| | 68 | 1 min (typically 1 min per kb DNA) |
| 1 | 68 | 1 min |
| 1 | 4 | ∞ |

#### Day 6-MB: Check your restreaked plates for isolated colonies

During the wait steps in today's protocols, check your restreaked plates from last lab period. If you still do not have isolated colonies, restreak your plate again, following the procedure from Day 5.

#### Day 6-PP: Specific Activity calculation (Described in Day 5-PP)

Remember that all units were calculated on Day 5?  there is time during this period to discuss any of the calculations and complete Table 4 if you did not do so during the last period. Show your results to teaching staff for confirmation.

### Day 7-PP: SDS-PAGE

#### Objectives

Today you will use sodium dodecyl sulfate polyacrylamide gel electrophoresis (SDS-PAGE) to separate your purification fractions and your cleared lysate by molecular weight. You will perform SDS-PAGE (on two separate gels) to assess the relative amount and purity of MDH in your large-scale induction cultures and subsequent purification fractions.

You will transfer the protein in one of these gels to nitrocellulose paper, a membrane used to immobilize protein. You will use this membrane for immunoblotting on Day 8. The second gel will be imaged via staining.

#### Background

##### Protein gel separations by SDS-PAGE

The primary uses for SDS-PAGE are the determination of the size and concentration of a protein, the estimation of protein purity in a solution, and fractionation of a protein mixture prior to immunoblotting. Fractionation of proteins using polyacrylamide gels is one of the principal means of protein characterization because of its speed and ease of use. The goal of SDS-PAGE is to separate proteins in a mixture solely based on molecular weight. The migration of a protein gives a good approximation of its size, and (after staining) the band intensity is a rough indicator of the amount of protein present in the sample.

Sample preparation is important for obtaining accurate separation of the proteins based on molecular weight. The sample treatments used should destroy any secondary or tertiary protein structure so these do not alter the migration of the proteins through the acrylamide matrix. Therefore, the sample preparation should solubilize and denature proteins, dissociate polypeptides, and reduce disulfide bonds. First, the protein sample is denatured with heat (boiling) in the presence of SDS and a reducing agent, usually dithiothreitol (DTT) or β-mercaptoethanol. SDS is an anionic detergent that denatures proteins by binding along polypeptide backbone. The SDS coats the proteins, providing them with a net negative charge proportional to their length. When treated with SDS, a reducing agent, and heat, the polypeptides become rods of negative charges with equal charge densities (charge per unit length). When the coated sample is run on an SDS-PAGE gel, the proteins separate by the sieving effect of the gel matrix.

Detection of proteins separated by SDS-PAGE will be accomplished using either Imperial or GelCode Blue stain. This Coomassie blue staining method is based on nonspecific (and irreversible) binding of Coomassie blue dye to proteins. The proteins are detected as blue bands of varying intensities on a clear background. One advantage of this method is that the fixation of the gel and proteins in acetic acid and methanol is eliminated, and often the gels do not need to be significantly destained prior to analysis.

#### Detection of a specific protein by immunoblotting

Like the Bradford assay, the Coomassie-based staining process is not specific for any particular protein. Because we are expressing MDH to a very high level and we know its molecular mass (~34kD when tagged), we may be able to putatively determine which band on the gel represents MDH in our purification fractions. To conclusively identify MDH subsequent to separation, we must use a technique specific for our protein. Immunoblotting (also called Western blotting) uses an antibody that specifically and with high affinity binds to our protein or to the affinity tag; the bound antibody can be readily detected to ultimately identify tagged MDH separated by the SDS-PAGE gel.

Immunoblotting is carried out in four steps: protein transfer to a solid support, blocking of unoccupied sites, primary and secondary antibody binding, and detection. After SDS-PAGE separation, the proteins must be transferred to a membrane since the gel will not allow ample diffusion of the antibody through the matrix to permit binding to the protein. Two different membranes are typically used- PVDF (polyvinylidene difluoride) and nitrocellulose. We use nitrocellulose in our lab due to its lower cost and suitability for routine immunoblotting procedures; PVDF is more hydrophobic and sturdy, and is typically used for applied techniques such as protein sequencing. The gel is pressed directly against the membrane, and proteins are transferred to the membrane using an electric current perpendicular to the gel. Transfers can be conducted in different formats (semi-dry or submerged), at different voltages (30V-100V), and for different lengths of time (one hour to overnight). Typically most if not all of the protein is transferred from the gel onto the membrane; the larger the protein and the higher the percentage gel, the more difficult transfer becomes. MDH is a moderately sized protein, and SDS-PAGE is performed with a 12.0% polyacrylamide gel to optimize transfer for immunoblotting.

Our lab will perform our transfers submerged in transfer buffer for one hour at 100V. Due to the high voltage and current, our transfer buffer will get quite hot and thus are performed on ice. Detection of MDH is then performed on the membrane. However, antibodies are polypeptides and can non-specifically associate with the membrane. After the proteins from the gel are transferred to the membrane, we must “block” the sites on the membrane that do not have any proteins transferred onto them. We will incubate our membrane in buffer containing milk protein (nonfat dry milk works well; purified casein can also be used) until the next lab period for this blocking step.

Antibodies with high affinity for the target protein must be used to allow for specific detection. Two classes of antibodies exist- monoclonal and polyclonal. Monoclonal antibodies are purified from a hybridoma culture expressing one antibody that recognizes exactly one part (epitope) of the protein (antigen). Polyclonal antibodies are isolated from serum of an animal injected with a pure protein, which in reality contains several antibodies each recognizing different epitopes of the same protein antigen. Monoclonal antibodies are less likely to be cross-reactive, and for identifying common protein targets are readily available (though more expensive). We will use a monoclonal antibody (produced in mouse) that recognizes the 6x His affinity tag.

Detection of the target protein occurs by detecting where on the membrane the antibody is bound. Most detection methods rely on conjugating the antibody to an enzyme, such as horseradish peroxidase (HRP) or alkaline phosphatase (AP). These enzymes catalyze reactions that either emit light (HRP) or form an insoluble precipitate on the membrane (AP) at the site of binding. However, coupling the antibody binding to the target protein may interfere with its binding affinity and is very expensive. Thus, conjugated secondary antibodies are employed to recognize the bound specific antibody (the "primary" antibody) at its constant region (which is determined by the host species producing the antibody). The advantages of a secondary antibody are three-fold: its affinity of binding is unaffected, signal amplification results since many secondary antibody molecules recognize one primary antibody, and only a handful of secondary antibodies are necessary to recognize all primary antibodies as only a few species of animals are used to produce primary antibodies.

#### Primary antibodies

Anti-6xHis - this means that it is an antibody that recognizes the His tag. This antibody requires the use of secondary anti-mouse AP-conjugated antibody. Detection is via a BCIP/NBT substrate, which forms an insoluble precipitate.

#### Procedure

Your group will be given two **pre**-cast 12% polyacrylamide gels to run your samples. Load each gel identically. Important safety note: unpolymerized acrylamide is a potent neurotoxin! Though these gels are pre-cast, there could be trace amounts of unpolymerized acrylamide present. Wear gloves at all times! These gels have 10 wells and one must contain a molecular weight marker (size standard). Therefore, you can load up to 9 samples onto each gel. You should run two identical gels. One will be used for antibody staining and one will be used for general protein staining. Each gel will be shared with one other group.

1. Prepare your pre-induced, cleared lysate, flow-through, and one elution. With the group you share your gel with, prepare one wash fraction. Mix 60 µL of each sample with 15 µL of 5X blue sample dye. Heat your samples for 5 minutes at 100ºC (heat block) to fully denature your proteins. Place samples immediately on ice until you are ready to load your gel.

2. Load 10 µL of the protein size marker (image of bands appears below) and 20 µL of each sample onto both gels. Add 20 µL of 1X loading dye in water to all empty wells. Note the exact locations of where your samples are loaded!!!

3. Make sure the center chamber is full with running buffer, to the line marked blotting (obtain help from teaching staff if needed), close the lid, and connect the leads to the power supply.

4. Run your gels at 200V until the blue dye front reaches the green gasket at the bottom of the gel (~45 min. running time)Turn off the power, remove the lid, and remove the gels from the plates with the help of the staff.

Note: one gel will be stained for 1 hour with Imperial Protein Stain (Thermo Scientific catalog #24615); the other will be used for immunoblotting. You will be working with both gels simultaneously. We recommend skipping to step 8 and returning to step 5 once transfer is started.

**Stain:**

5. Rinse gel with distilled water and leave in room temperature distilled water for 5 min.

6. Remove water, add ~15 mL Imperial Protein Stain or Acqua Stain if provided, and leave on shaking platform at room temperature overnight. The teaching staff will pour off the stain and add water to destain the gel until the next lab period.

7. Image gel using gel documentation system (make sure to get a high quality picture, with no extraneous light effects and good focused bands) and analyze. Label each molecular weight marker and indicate on the gel the band(s) you are putatively identifying as MDH. Can you determine visually any tagged and untagged MDH? Estimate the molecular weight of MDH by generating a molecular weight standard curve using the size markers and migration distance(s) (plot log MW vs. migration distance in mm). If you are unable to identify the MDH band(s), you may analyze your immunoblot for size determination of MDH.

**Blot:**

8. Prepare the other gel for transfer. Make sure you have one nitrocellulose membrane and two pieces of filter paper cut to the same size as your gel before proceeding. Important: do not touch the membrane with bare hands, or proteins in your hands will stick to the membrane! Handle at the corners with forceps!

9. The apparatus will be assembled as shown in the figure 6 below. It is critical to assemble as depicted, so that the gel and membrane are properly oriented with respect to the current!

Figure 8: Western Blot precast polyacrylamide gel, membrane and accessories set up

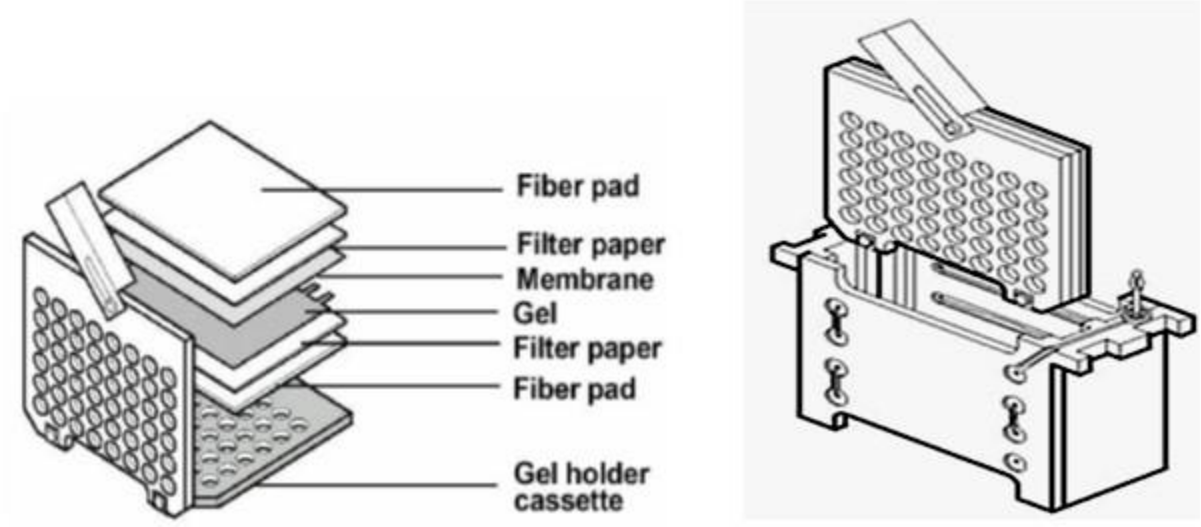

Western Blot precast polyacrylamide gel, membrane and accessories set up (left) and complete electrode assembly unit (right)

10. Fill a small container halfway with transfer buffer.  Take two of the fiber pad sponges and submerge in buffer.  Be sure to press the sponges into the solution to fully hydrate them - air bubbles that remain will inhibit transfer of proteins!

11. In another container filled 1/3 with transfer buffer, pre-wet the filter paper sheets and membrane briefly using forceps.  Assemble the transfer apparatus as demonstrated and shown in the figure above.

12. Insert an ice brick, fill apparatus tank with transfer buffer, and put the tank on ice (use an ice bucket or blue bucket).

13. Connect the lid and leads, and transfer at 100V for one hour.  Turn off power supply, disconnect, and remove membrane from transfer sandwich with forceps and place in a clean container.

14. Add PBS-T to the membrane so that the membrane is entirely submerged, cover the container, and incubate at 4ºC until the next lab period.

Note:  Markers used are the EZ-Run Pre-stained Rec Protein Ladder, and an image of what these markers will look like on your gel appears in figure 7 below.  Information that follows is taken from the Fisher Scientific website (fishersci.com).

Figure 9: Bands showing the Fisher BioReagents* EZ-Run* Prestained Rec Protein Ladder

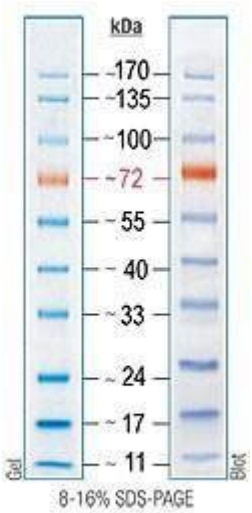

This marker is comprised of ten recombinant proteins covalently coupled to a blue chromophore with the 72kDa reference band tagged with orange dye. The 24 kD band and all lower molecular weight bands may not be visible using our gel system.

### Day 8-PP: Enzyme kinetics

#### *Objectives*

This week we will test the function of our purified MDH enzyme in vitro using enzyme kinetics: the study of enzyme rates. We will be using variations on the MDH activity assay we used during purification to calculate $V_{max}$ and $K_m$. This requires determining the effect of substrate concentration [S] on the rate of hydrolysis (velocity of the reaction) and using these data to determine the maximal velocity for the enzyme at a given substrate concentration ($V_{max}$), as well as the substrate concentration (Km) at which the enzyme has achieved half the $V_{max}$. You will be able to compare two methods of graphing enzyme kinetics data: Michaelis-Menten (M-M) and Lineweaver-Burk (L-B) analyses.

#### *Background*

##### Why study enzyme kinetics?

Determination of kinetic parameters and understanding enzyme saturation behavior are important for understanding enzyme properties. For example, lactate dehydrogenase (LDH) activity appears in blood plasma only after certain cells are damaged, allowing the enzyme to leak out. Because the kinetic characteristics of heart muscle LDH are different from those of isoenzymes in other tissues, the kinetic analysis of blood can be used to measure damage caused by myocardial infarction (heart attack). Detailed kinetic analyses of enzymes are additionally important in drug design and in determining the dosage appropriate for the treatment of a variety of diseases.

##### Drugs that act as specific enzyme inhibitors

Many commonly used medicines act as inhibitors of enzymes. Many fungi and plants naturally produce antibiotic compounds to kill bacteria in their environment. These compounds inhibit enzymes responsible for cell wall synthesis (penicillin), DNA replication (Cipro®), or protein synthesis (erythromycin) in pathogenic bacteria but not analogous enzymes in mammalian cells.

Biochemists have studied natural inhibitors and have chemically synthesized related compounds to determine the kinetic parameters such as Km and Vmax of enzymes *in vitro*. By understanding how enzymes function, one can design new inhibitor compounds with specific biomedical or agricultural applications (typically referred to as rational drug design). Rational drug design is a burgeoning field in which detailed information about an enzyme's structure and kinetic parameters are used to design inhibitor compounds that can be synthesized and put to good use. Tamiflu® (oseltamivir) is synthesized by Roche Pharmaceuticals (roche.com) and is currently used at the first signs of flu-like symptoms (or even prophylactically) to decrease the severity and duration of influenza. Flu virus harbors a neuramidase enzyme, which is required for viral release from infected host cells. Oseltamivir competitively inhibits the viral neuramidase enzyme from cleaving the virus bound to sialic acid linked to host cell surface proteins. Additional examples of inhibitors synthesized in the laboratory are the anti-viral drug AZT and the herbicide ROUNDUP™.

AZT is a competitive inhibitor of viral (but not human) DNA polymerase and is widely used to combat HIV infection. ROUNDUP™ is a supposedly safe and effective herbicide because it functions as a competitive inhibitor of an aromatic amino acid biosynthetic enzyme found only in plants and microbes.

#### Kinetics and calculation of $K_m$ and $V_{max}$

The product formation rate is dependent on the intrinsic kinetic parameters ($V_{max}$ and $K_m$). Recall that enzyme kinetics can often be described using the Michaelis-Menten relationship (see equation 3) to determine the reaction rate. This relationship was derived based on the fact that enzyme and substrate achieve a rapid equilibrium to form the enzyme-substrate complex, which then dissociates to give product and free enzyme (see equation 1). This relationship can also be achieved based on the quasi-steady state assumption proposed by Briggs and Haldane, which assumes that the concentration of the enzyme-substrate complex is essentially constant; however, this assumption fails in the early stages of the reaction. The derivation of the Michaelis-Menten equation starts by writing the first order kinetic equation for change in product over time (see equation 2).

$$E + S \underset{k_{-1}}{\overset{k_1}{\rightleftharpoons}} ES \xrightarrow{k_2} E + P \qquad \text{(eq. 1)}$$

where E is enzyme, S is substrate, ES is the enzyme-substrate complex, P is product, E is free enzyme, and $k_1$, $k_{-1}$ and $k_2$ are reaction rate constants.

$$v = \frac{dP}{dt} = k_2[ES] \qquad \text{(eq. 2)}$$

After substituting for [ES] and using an enzyme mass balance where $[E_0] = [E] + [ES]$, the final form of the Michaelis-Menten equation becomes:

$$v = \frac{v_{max}[S]}{[S] + K_m} \qquad \text{(eq. 3)}$$

where $v_{max} = k_2[E_0]$ and $K_m = \frac{k_{-1} + k_2}{k_1}$

This equation can be rearranged to yield the Lineweaver-Burk equation to linearize the data:

$$\frac{1}{v} = \frac{1}{v_{max}} + \frac{K_m}{v_{max}}\frac{1}{[S]} \qquad \text{(eq. 4)}$$

We can effectively design experiments to calculate both $V_{max}$ and $K_m$ for an enzyme and its reaction with a specific substrate. You will need to determine the reaction velocity (v) at different substrate concentrations [S]. Once these data have been collected, both $V_{max}$ and $K_m$ can be determined by either plotting v vs [S] in a traditional Michaelis-Menten saturation curve or using the Lineweaver-Burk double reciprocal rearrangement of the data.

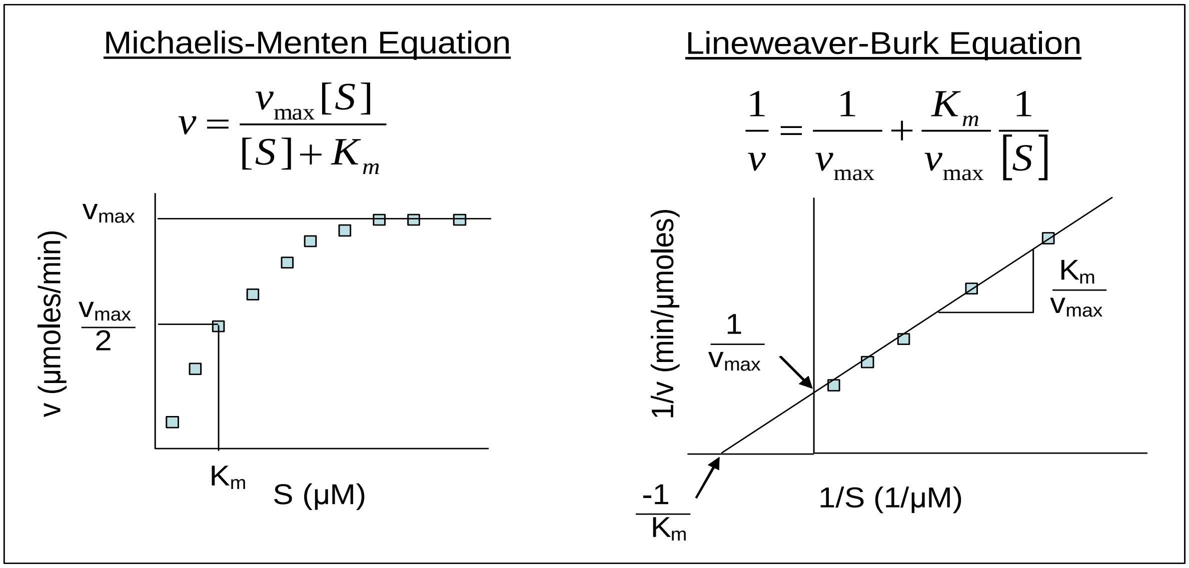

Figure 10. Examples of the equations, and resulting plots for, Michaelis-Menten equation (left) and Lineweaver Burk equation (right).

#### Enzyme Inhibition.

Reversible inhibition (one in which the inhibitor can bind to and unbind from the enzyme) is classically divided into categories (competitive and noncompetitive for this laboratory). Competitive inhibitors resemble the substrate, and thus compete with the substrate for binding to the active site. The inhibitor-enzyme complex precludes binding of substrate. Since the inhibitor binding is reversible (not permanent), once the inhibitor dissociates from the enzyme, substrate can bind. The degree of inhibition of the enzyme activity depends upon the dissociation rate of the inhibitor. Competitive inhibitors often have a higher binding affinity for the enzyme than the substrate does, so if there is a mixture of substrate and inhibitor, inhibitor will bind more often and less product will be made. Increasing the substrate concentration can overcome the effects of the competitive inhibitor. In kinetic terms competitive inhibition is manifested in a change in the Km . Recall that Km is the substrate concentration at which the enzyme attains half its maximal velocity. Competitive inhibitors raise the Km because it takes higher concentrations of substrate to out-compete the inhibitor. Eventually you could add enough substrate to overcome inhibition; therefore, the Vmax is unchanged.

**Noncompetitive inhibitors** do not bind the enzyme in the same place as the substrate, so both inhibitor and substrate can bind. Upon binding, these compounds alter the conformation of the enzyme (the three-dimensional structure) so the enzyme can no longer catalyze the chemical conversion of the substrate. The conformation of the active site, which is separate from the inhibitor-binding site, is altered and rendered non-functional. With non-competitive inhibitors, increasing the substrate concentration has no effect on inhibition, since substrate and inhibitor are not in competition for the same binding site. In kinetic terms, noncompetitive inhibition is manifested in a lowering of $V_{max}$, but the $K_m$ stays the same. Lineweaver-Burk plots of competitive and non-competitive inhibition are shown below.

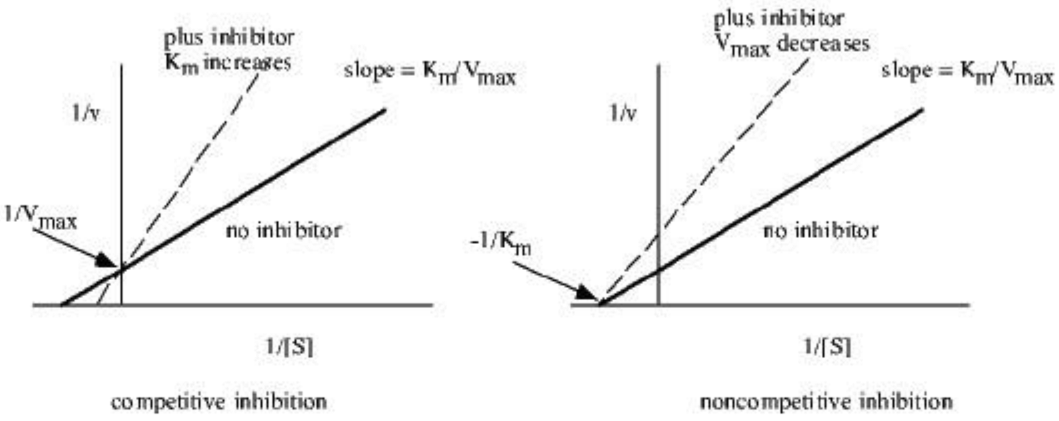

#### Quantitation of MDH activity

Recall that MDH catalyzes the reversible reaction Malate → Oxaloacetate (OAA), which reduces the cofactor NAD+ to NADH.

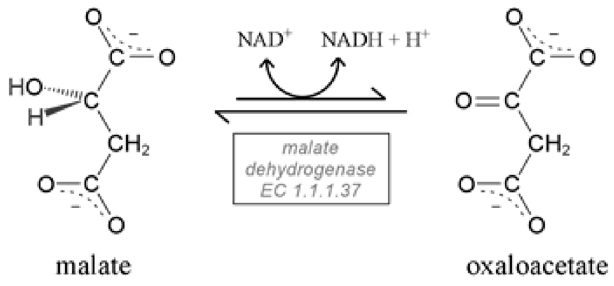

For MDH assays, we monitor the reverse reaction: Oxaloacetate → Malate and take advantage of the absorbance at 340nm of NADH. Loss of absorbance at 340 nm represents the conversion of NADH to NAD+. We can convert change in absorbance per min into units per ml of MDH activity.

To determine $K_m$, we will *vary the concentration of oxaloacetate*, but be sure to keep the concentration of NADH constant (and high enough that it is not limiting for the reaction to proceed.

#### Procedures

*****Measuring Michaelis Menten (M-M) parameters takes time, flexibility and patience*****

##### Experiment I: Michaelis-Menten Curve with purified MDH.

Your primary goal today is to produce a good Michaelis-Menten curve of your purified MDH. Once this is achieved, you can set up Experiment II.

##### Experiment II: Vary conditions that affect the Michaelis-Menten parameters of MDH.

Once you have good data for Experiment I, you can ask teaching staff about doing a comparison in which you vary one of the conditions of the reaction.

##### The MDH Assay

As you have done before, you mix substrate (OAA) and NADH in assay buffer (10mM K/Na Phosphate Buffer, pH 7.4), add your enzyme sample and allow the reaction to proceed for a specified time, at which point the reaction is stopped by adding a 1 M sodium bicarbonate. The absorbance at 340nm is determined (compared to a control reference sample that contains everything but enzyme and thus will have no conversion of NADH).

The curve is generated by plotting $v_o$ against OAA concentration.

##### Determining $v_o$:

The time your reactions run needs to be short enough that product formation is in the linear range (i.e. reactions have not begun to run out of substrate or cofactor, or begun to slow down for any other reason). You might use the same amount of time you used previously, but you do not want your reactions to have run to completion for this experiment, so you might choose a shorter time period. This is known as the initial (linear) velocity, also known as $v_{o.}$

This velocity is a measure of the rate of change of NADH concentration, which is calculated from absorbance, factoring in the pathlength of sample liquid (L) and extinction coefficient for NADH ($\varepsilon = 6200\ M^{-1}cm^{-1}$). In a microplate 100 µL gives L = 0.28cm.

$$\text{Reaction velocity } v\ (\mu\text{mol NADH/min}) = \frac{(\Delta A_{340})\ (0.0001\text{L})\ \ (10^{6}\ \mu\text{mol})}{(6200\ 1/\text{M}\ 1/\text{cm})(0.28\text{cm})(\text{assay time in min.})\ (\text{mol})}$$

By combining constants:

$$v = \frac{(\Delta A_{340})\ \ (0.058\ \mu\text{mol})}{(\text{time in min.})}$$

USE THIS EQUATION FOR YOUR CALCULATIONS!

#### Experiment II: Michaelis-Menten Curve with purified MDH: reaction velocity V substrate concentration

Preparing enzyme reactions:

*Components:*
**Assay Buffer**: 10 mM K/Na Phosphate Buffer, pH 7.4
Keep at room temperature.

**NADH**: Provided at **6** mM. Use at a final concentration of **0.6** mM.
Keep on ice.

**Enzyme:** *Keep on ice at all times!*
You have calculated the concentration of your MDH protein solution using the Bradford Protein assay. Determine the working concentrations of MDH stock that you want to use. You will probably use a concentration similar to the one that worked when you were calculating specific activity, however if you very high activity (completely consumed NADH), or low activity, you might need to adjust this concentration). Consult with the teaching staff about the concentration you have chosen.

**Substrate [S]**: oxaloacetate (OAA) provided at 20 mM in assay buffer (Keep on ice)

. You want to add different amounts of this stock to give you a range of (final) concentrations: the range should be within **0.05-2** mM final concentration of OAA.

**STOP Solution:** 1 M $Na_2CO_3$ (Keep at room temperature)

##### Setting up dilutions of OAA:

You probably should not expect to be able to accurately pipette volumes of less than 2 uL into your reaction tubes, so you should make appropriate dilutions of your OAA stock in order to get good values for concentrations in the lower range.

For example: Make a 1/10 dilution of OAA and use 30 uL in a reaction for a final concentration of 0.2mM OAA, or 15 uL for a final concentration of 0.1 mM OAA.
Determine the amounts to add to each tube by completing Table 7: Worksheet

##### Plan your reactions:

Length of time chosen for reaction: ____________________

Table 7: Worksheet:

You may use the table below to calculate amounts to use.
Items in brackets "[ ] " indicate target final concentrations. Some sample concentrations are included as a reference (although you do not need to use these examples). Most of you will start with 7-10 different [S], there are more rows in this table to allows for additional reactions.

| **Rxn #** | **NADH (μL)** | [MDH] (μg/mL) | **Vol. MDH (μL)** | [OAA] (mM) | **Vol. OAA (μL)** | **Buffer (μL)** | **Total Vol. So far (μL)** | **Stop (μL)** |
|---|---|---|---|---|---|---|---|---|
| **Ctrl 1** | **0** | | | | | | **300** | **30** |
| **Ctrl 2** | **30** | -- | **0** | | | | **300** | **30** |
| **3** | **30** | | | 2mM | 30ul N | | **300** | **30** |
| **4** | **30** | | | 1mM | 15ul N | | **300** | **30** |
| **5** | **30** | | | 0.75mM | 11.25 N | | **300** | **30** |
| **6** | **30** | | | | | | **300** | **30** |
| **7** | **30** | | | 0.2mM | 30ul 1/10 | | **300** | **30** |
| **8** | **30** | | | 0.1mM | 15ul 1/10 | | **300** | **30** |
| **9** | **30** | | | 0.075mM | 11.25 1/10 | | **300** | **30** |
| **10** | **30** | | | | | | **300** | **30** |
| **11** | **30** | | | | | | **300** | **30** |
| **12** | **30** | | | | | | **300** | **30** |
| **13** | **30** | | | | | | **300** | **30** |
| **14** | **30** | | | | | | **300** | **30** |
| **15** | **30** | | | | | | **300** | **30** |
| **16** | **30** | | | | | | **300** | **30** |
| **17** | **30** | | | | | | **300** | **30** |
| **18** | **30** | | | | | | **300** | **30** |
| | **30** | | | | | | **300** | **30** |
| | **30** | | | | | | **300** | **30** |
| | **30** | | | | | | **300** | **30** |
| | **30** | | | | | | **300** | **30** |
| | **30** | | | | | | **300** | **30** |
| | **30** | | | | | | **300** | **30** |
| | **30** | | | | | | **300** | **30** |
| | **30** | | | | | | **300** | **30** |
| | **30** | | | | | | **300** | **30** |
| | **30** | | | | | | **300** | **30** |
| | **30** | | | | | | **300** | **30** |
| | **30** | | | | | | **300** | **30** |
| | **30** | | | | | | **300** | **30** |
| | **30** | | | | | | **300** | **30** |
| | **30** | | | | | | **300** | **30** |

- 

- For help with calculations:

$M_1V_1 = M_2V_2$
Solve for $V_1$
$M_1$ = stock concentration (6mM NADH, etc.) $M_2$ = final conc (0.6mM)
$V_2$ = final vol 300 µL

#### Turnover number

In enzymology, the turnover number ($k_2$) of an enzyme (in our case MDH) is defined as the number of substrate molecules (in our case oxaloacetate) converted into product by an enzyme molecule in a unit time when the enzyme is fully saturated with the substrate (i.e., when every catalytic site is occupied). Alternatively, this can be written as:

$$k_2 = \frac{v_{max}}{E_t} \quad (5)$$

Where $k_2$ is the turnover number ($min^{-1}$) and $E_t$ is the amount of active sites (moles). (moles/min) is determined as above and can be estimated as described below:

$$E_t = [MDH]_{RXN} / MW_{MDH} \times V_{RXN} \quad (6)$$

is the total volume of the reaction mixture. The molecular weight of MDH in kDa (1 Da = 1 g/mol) is known for our tagged enzyme and can also be estimated from SDS-PAGE. The concentration (mg/mL) of MDH in the sample that you use in the kinetics assays can be determined from the following relation:

$$[MDH]_{sample} = [total\ protein]_{fraction} \times purity \quad (7)$$

The total protein (mg/mL) in that sample is estimated using the Bradford assay. The purity can be estimated from the SDS-PAGE Once you have calculated the concentration of MDH (mg/mL) in the sample or fraction, you can determine the
concentration of MDH used in the reaction mixture from the following relation:

$$[MDH]_{RXN} = [MDH]_{fraction} \times V_{fraction\ used\ in\ assay} / V_{RXN} \quad (8)$$

Once the concentration of MDH in your reaction is known, you can determine from equation 6 and the turnover number from equation 5. The turnover number is a commonly used indicator of enzyme efficiency.

Information you will need: Size of protein
Estimated purity = Concentration of protein

#### Procedure

##### Running the Reactions

*As a first run, setup 7-10 reactions with different [S] concentrations and your wild type enzyme to make sure the experiment works well:*

*Determine a reasonable interval to stagger your timed reactions so that you have time to start and stop successive reactions in a timely manner*

1. Set up micro centrifuge tubes with volumes as directed in your completed table.

- Add Buffer: One option for ease, add 200 uL buffer to each tube first, then go through and add the remaining amount (generally 40-70 uL) as appropriate to each of the tubes.

- Add NADH and OAA. Let these solutions get to room temperature ( a few minutes)

- DO NOT ADD ENZYME YET!!

2. Do not forget the critical controls –

   1) Enzyme, assay buffer and OAA (highest conc.), **but <u>no NADH</u>**
   2) NADH, assay buffer and OAA (highest conc.), **but <u>no enzyme</u>**

3. Start the reaction by adding enzyme to the appropriate tube

4. Vortex briefly and incubate at room temperature for two minutes.

5. Add 30 μL of STOP solution and vortex immediately to each tube in order to stop the reaction (this will change the final volume of reaction, but we will can ignore this change for purposes of calculation of v).

6. Transfer 100 μL into each of two wells of a microplate (each of your reactions should be read in duplicate).

7. Read the absorbance at 340 nm and record in your table (remember to record the Δ absorbance).

8. Determine reaction velocities (see **Table 2: Results**)

<u>Now: Determine if this experiment worked</u>: did you get a loss in absorbance in the tubes you expected? Is there a difference at different substrate concentrations? If not, you might need to vary the [MDH], the [S] or the time of reaction.

Plot your M-M curve (instructions below)- see if the data look good before proceeding (get someone from the teaching staff to check it for you)

a) If yes- do you need more [OAA] concentrations to have a nice curve?
b) If no, discuss plans, change some variables: Continue until data look good

IMPORTANT: Your data will be drastically affected by pipetting errors. If your duplicates are not close in value, or your data points are wildly different from the expected values, discuss this with the TAs or instructor. Also, keep in mind that the **data points for the lowest change in absorbance (at the lowest substrate concentrations) are the least reliable**. It may be necessary to ignore the data for certain plots.

If all is working well, complete the other reactions that you need, i.e. other concentrations of [S] to complete Experiment 1. Then prepare for Experiment II.

Table 8: **Results:** table for calculation of reaction velocities.

| **Rxn #** | Control 2 | Sample Abs. | Δ Abs. | Constant (μmol) | Time (min) | **v (μmol NADH/min)** |
|---|---|---|---|---|---|---|
| **3** | | | | 0.058 | | |
| **4** | | | | 0.058 | | |
| **5** | | | | 0.058 | | |
| **6** | | | | 0.058 | | |
| **7** | | | | 0.058 | | |
| **8** | | | | 0.058 | | |
| **9** | | | | 0.058 | | |
| **10** | | | | 0.058 | | |
| **11** | | | | 0.058 | | |
| **12** | | | | 0.058 | | |
| **13** | | | | 0.058 | | |
| **14** | | | | 0.058 | | |
| **15** | | | | 0.058 | | |
| **16** | | | | 0.058 | | |
| **17** | | | | 0.058 | | |
| **18** | | | | 0.058 | | |
| | | | | 0.058 | | |
| | | | | 0.058 | | |
| | | | | 0.058 | | |
| | | | | 0.058 | | |
| | | | | 0.058 | | |
| | | | | 0.058 | | |
| | | | | 0.058 | | |
| | | | | 0.058 | | |
| | | | | 0.058 | | |
| | | | | 0.058 | | |
| | | | | 0.058 | | |
| | | | | 0.058 | | |
| | | | | 0.058 | | |
| | | | | 0.058 | | |
| | | | | 0.058 | | |
| | | | | 0.058 | | |
| | | | | 0.058 | | |

#### Construct Michaelis-Menten (M-M) and Lineweaver-Burk (L-B) plots:

To calculate $K_m$ and $V_{max}$ for the enzyme. You will consider which you think is a better approach with your data.

#### Using Excel to Plot Enzyme Kinetics and Inhibition Data:

Open Excel. Click on the top left spreadsheet box and type in "**[S]**". This will be your **X-axis** for the Michaelis-Menten plot. Enter the values for substrate concentrations. Click on the first row cell in column B and type in "**v**". This is the **Y-axis** for your Michaelis-Menten plot. Enter the values for velocity that you calculated. Continue to enter 1/[S], 1/v, etc.

#### The Michaelis-Menten (M-M) Plot

Highlight the data in columns A and B. Click the Insert tab, then on the Scatter tab, and select the scatter plot style without any lines. To properly format your graph, click on the Layout tab to add titles. Use an appropriate chart title (such as Michaelis-Menten), label the axes (do not forget units!) and remove the legend. Change other plot parameters as you wish (most are handled under the Design tab). Do not add a trendline! Estimate $V_{max}$ and $K_M$ from plot.

#### The Lineweaver-Burk (L-B) Plot:

Go back to your table on the worksheet and select the data in the 1/v and 1/[S] columns for L-B. Graph these data points as above, using a scatter plot as you did for the M-M plot. Now you need to generate a line and the equation that describes the relationships. Be sure to have the chart you are working on selected. Under the Layout tab, click on Trend line to add the line of best fit. You will perform a linear fit, and be sure to add the equation of the line and R-squared value on the plot (this is under "more trend line options"). Determine $K_m$ and $V_{max}$ from the linear equation that describes your data.

**MDH**

| **Graphing method** | **$K_m$ (mM)** | **$V_{max}$ (µmol/min)** |
|---|---|---|
| **Michaelis-Menten** | | |
| **Lineweaver-Burk** | | |

**Show your** Michaelis Menten curves from Experiment I to the teaching staff to confirm that you have a sufficient result.

### Experiment II: Vary conditions that affect the Michaelis-Menten parameters of MDH

In these studies, you will be collecting kinetic measurements by varying some condition of the reactions. This could include a change in temperature (10C-60C), change in pH (pH2-10), or addition of inhibitor (Choose an inhibitor: citrate, succinate, alpha-ketoglutarate, alpha-ketobutyrate, or pyruvate, supplied at 500 mM).

You will set up this experiment as you did Experiment I, but with this variation in conditions.

Table 9: Worksheet for Experiment II

| Rxn # | NADH (µL) | [MDH] (µg/mL) | Vol. MDH (µL) | [OAA] (mM) | Vol. OAA (µL) | Buffer (µL) | Total Vol. So far (µL) | Stop (µL) |
|---|---|---|---|---|---|---|---|---|
| Ctrl 1 | 0 | | | | | | 300 | 30 |
| Ctrl 2 | 30 | -- | 0 | | | | 300 | 30 |
| 3 | 30 | | | | | | 300 | 30 |
| 4 | 30 | | | | | | 300 | 30 |
| 5 | 30 | | | | | | 300 | 30 |
| 6 | 30 | | | | | | 300 | 30 |
| 7 | 30 | | | | | | 300 | 30 |
| 8 | 30 | | | | | | 300 | 30 |
| 9 | 30 | | | | | | 300 | 30 |
| 10 | 30 | | | | | | 300 | 30 |
| 11 | 30 | | | | | | 300 | 30 |
| 12 | 30 | | | | | | 300 | 30 |
| 13 | 30 | | | | | | 300 | 30 |
| 14 | 30 | | | | | | 300 | 30 |
| 15 | 30 | | | | | | 300 | 30 |

MDH, Experiment II.

| Graphing method | Km (mM) | Vmax (umol/min) |
|---|---|---|
| Michaelis-Menten | | |
| Lineweaver-Burk | | |

#### Discussion Questions:

Compare M-M and L-B graphing methods for determining kinetic parameters. Do they yield similar results? Which seems to have more likely error in the calculation? What approach might have the most error if there are few time-points? What approach might have the most error for the samples at lowest OAA concentration?

Compare results for the two experiments (I and II). Are the results different? What is the likely reason for any variation? Can you explain why you think the parameters changed the way they did?
