## Supplementary document 2 for "CUR(E)ating a New Approach to Study Fungal Effectors and Enhance Undergraduate Education through Authentic Research"

Supplementary document B

Sample Schedule:

Gateway cloning[1]: Gateway™ LR Clonase™ II Enzyme mix was purchased from Invitrogen (Waltham, MA). The protocol was adapted from the manual: Make a 8 μL solution containing: 50-150 ng of entry vector and 150 ng of destination vector. Add 2 μL of the LR Clonase™ II enzyme mix, briefly vortex twice, and incubate the LR mixture at 25°C for 18 hours. Then, 1 μL of Proteinase K is added to the mixture, incubate at 37°C for 10 minutes, and 2 μL of the LR reaction product will be used for transformation.

Agro-transformation: GV3101 ElectroCompetent Agrobacterium was purchased from Intact Genomics (St. Louis, MO) and protocol was adapted from the manual: 1 μL of 100 ng plasmid DNA is added to 25 μL of competent cells, mixed by gentle tapping, and transferred to a pre-chilled, 1 mm electroporation cuvette without introducing bubbles. Electroporate the cell mixture at 1800 V, immediately resuspend the cells with 974 μL of recovery medium, transfer to a microcentrifuge tube, and incubate at 30°C, 200 rpm for 3 hours. Then the cell mixture can be diluted and spread onto plates containing proper selective marker.

Agro-infiltration[2]: Protocol for agro-infiltration was adapted from Goodin, et al. 2002 with minor changes: Inoculate single colony or glycerol stock of *A. tumefaciens* transformant culture in 5 ml LB with selective marker. On the second day, inoculate 20 μL of the overnight culture into fresh 50 mL culture with selective marker. On the next day, measure the OD_600_, dilute, and resuspend the cells into OD_600_ = 0.6 with MMA buffer (200 μM acetosyringone, 10 mM MgCl_2_, 10 mM MES, pH5.6). Use a 1 mL needleless syringe to infiltrate the abaxial surface of a 6-week-old *N. tabacum* leaf. Tissues will be ready for microscope analysis at 3 days post infiltration.
